# Long-read Transcriptome Landscapes of Primary and Metastatic Liver Cancers at Transcript Resolution

**DOI:** 10.1101/2023.07.11.548526

**Authors:** Zhiao Chen, Qili Shi, Yiming Zhao, Midie Xu, Yizhe Liu, Xinrong Li, Li Liu, Menghong Sun, Xiaohua Wu, Zhimin Shao, Ye Xu, Lu Wang, Xianghuo He

## Abstract

**Background:** The liver is the sixth most common site of primary cancer in humans and is frequently colonized by metastases from cancers of other organs. Few studies have investigated the transcriptomic profiles of matched primary tumor and hepatic metastases of patients. Moreover, the read length of 100-200 bases in conventional short-read RNA sequencing is too short, which makes it difficult to directly infer the full-length transcript structure. To help develop effective treatments and improve survival, it is crucial to understand the complex and diverse molecular mechanisms of primary and metastatic liver cancers.

**Methods:** Ninety-five primary and secondary liver cancer patients who underwent hepatic resection were included with long-read sequencing isoform-sequencing and short-read RNA sequencing. We compared the transcriptome landscapes of primary and metastatic liver cancers and systematically investigated HCC, paired primary tumors and liver metastases, and matched non-tumor liver tissues.

**Results:** We defined the full-length isoform-level transcriptome of human primary and metastatic liver cancers and identified isoform-level diversity in HCC and metastasis-associated transcriptome variations in metastatic liver cancers. Specific RNA transcripts and isoform switching events with clinical implications were profoundly discovered in liver cancer. Metastasis-specific transcripts that can predict the metastatic risk and identify the primary sites of cancers of unknown primary liver metastasis patients were defined. Additionally, we found that adjacent paracancerous liver tissues are abnormal and characterized the premetastatic immunological and metabolic alterations in the liver that favor the spread of cancer metastases.

**Conclusions:** Our findings strongly highlight the powerfulness of full-length transcriptome profiling to yield novel biological insights into understanding the molecular basis of tumorigenesis and will further benefit the treatment of primary and metastatic liver cancers.

## Background

Liver cancer is the fourth leading cause of cancer-related mortality worldwide. Hepatocellular carcinoma (HCC) accounts for 80-90% of primary liver cancers, and generally arises in a background of cirrhosis and inflammation[1]. Liver metastases are commonly detected in a range of malignancies, including colorectal cancer (CRC), pancreatic cancer, melanoma, lung cancer and breast cancer, although CRC is the most common primary cancer that metastasizes to the liver[2]. In general, the occurrence of liver metastases is significantly correlated with a decrease in the 5-year survival rate and reduced quality of life[3]. Although surgical resection has largely been used to treat well-selected patients with liver metastasis, the prognosis of liver metastases is poor[4]. Large-scale cancer sequencing has focused on primary tumors, such as HCC, and identified various molecules that regulate this complex multistep process, highlighting the complex molecular and cellular changes that occur in liver cancer[5, 6]. Dysregulated processes such as epithelial to mesenchymal transition (EMT), morphological pattern formation, cancer stem cell regulation, and microenvironment remodeling are critical for tumor cells to acquire metastatic capacity[7]. Cancer metastasis is a process involving local invasion, intravasation, survival in the blood circulation, extravasation, adaptation to survival in a new microenvironment, and colonization and outgrowth in distant body sites[8]. However, few studies have investigated the transcriptomic profiles of matched primary tumor and hepatic metastases of patients. Moreover, the read length of 100-200 bases in conventional short-read RNA sequencing is too short, which makes it difficult to directly infer the full-length transcript structure[9]. To help develop effective treatments and improve survival, it is crucial to understand the complex and diverse molecular mechanisms of primary and metastatic liver cancers.

Long-read sequencing allows the comprehensive analysis of transcriptomes by identifying full-length splice isoforms and several other posttranscriptional events[10, 11]. Analysis of eight patient-derived HCC cases and the Hep3B cell line by long-read SMRT sequencing revealed that alternative isoforms (AS) and tumor-specific isoforms that arise from aberrant splicing are common during liver tumorigenesis[12]. Another study used Nanopore RNA-seq on three HCC patients’ tumors, matched portal vein tumor thrombi, and peritumoral tissues and discovered two novel prognostic transcripts[13]. A recent study developed an analysis pipeline with an Oxford Nanopore sequencer and revealed novel splicing abnormalities and oncogenic transcripts in liver cancer[14]. Although these studies have demonstrated that AS of mRNA precursors plays important roles in HCC development, it is difficult to illustrate the whole AS landscape of HCC, and the full-length transcriptomes of primary and metastatic liver cancers have remained underexplored.

Accurately quantifying transcripts using long-read sequencing requires the depth of coverage, which is prohibitively expensive. Therefore, in this study, we combined long-read sequencing isoform-sequencing (Iso-Seq) with short-read RNA sequencing to survey the transcriptome landscapes of primary and metastatic liver cancers and systematically investigated HCC, paired primary tumors (PTs) and liver metastases (LMs), and matched non-tumor liver tissues (LM-NTs). Our results provide the first full-length primary and metastatic liver cancer transcriptome profiles at transcript resolution, which would be useful to understand the molecular basis of liver cancer and will further transform the approach to treating primary and secondary liver cancers.

## Methods

### Patients and sample characteristics

We collected samples of pathologically diagnosed liver metastases from Fudan University Shanghai Cancer Center with written informed consent and approval from the Ethics Committee of the Fudan University Shanghai Cancer Center (approval number: 2011-ZZK-33). The study was conducted in accordance with ethical guidelines (Declaration of Helsinki). In detail, 177 samples from 95 patients (n = 23, HCC; n = 7, nasopharyngeal carcinoma; n = 12, breast cancer; n = 5, gastric cancer; n = 3, kidney cancer; n = 5, neuroendocrine tumor; n = 20, colorectal cancer; n = 18, ovarian cancer; n = 2, cervical cancer) underwent Iso-seq, and 203 samples from these patients were performed by RNA-seq. Another 62 liver tissues from 62 patients were analyzed by IHC (n = 2, nasopharyngeal carcinoma; n = 6, breast cancer; n = 1, kidney cancer; n = 20, colorectal cancer; n = 11, ovarian cancer; n = 3, cervical cancer; n = 19, hepatic hemangioma).

### Statistical analysis

Wilcoxon rank-sum test was used to determine the statistical difference between two groups. Multiple testing corrections were performed with the Benjamini and Hochberg method. Kaplan–Meier survival curves were generated by using the R package “survminer” (version 0.4.8). Log-rank tests were utilized to compare the overall survival difference in groups by using the R package “survival” (version 3.2–3). The chi-squared (χ2) test was used to evaluate the different expression of PD-1 and PD-L1 between liver tissues from LM patients and hemangioma patients. Statistical analyses were performed by SPSS (IBM, NY, USA) and R project. A p-value < 0.05 was considered statistically significant.

**The details of Methods are provided in Additional file 1: Supplementary Methods.**

## Results

### Long-read transcriptome landscapes of primary and metastatic liver cancers

Ninety-five primary and secondary liver cancer patients who underwent hepatic resection were enrolled in this study. Regarding patient demographics, there were 41 (43.2%) males and 54 females (56.8%). The median age of the cohort was 55 years, with a range of 31-77 years **(Additional file 2: Table S1)**. **Figure 1a** shows the cancer types represented in the cohort. The top three liver metastases cancer types in our cohort included colorectal cancer liver metastases (CRLM), breast cancer liver metastases (BCLM), and ovarian cancer liver metastases (OCLM). We obtained 203 tissues from these patients, including paired HCC and adjacent non-tumor tissues (HCC-NT), paired primary tumors (PT) and liver metastases (LM), and matched non-tumor liver tissues (LM-NT). A total of 203 and 177 tissues were successfully evaluated by short-read sequencing and Iso-Seq, respectively. Detailed information is listed in **Additional file 2: Table S1.**

**Figure 1.**
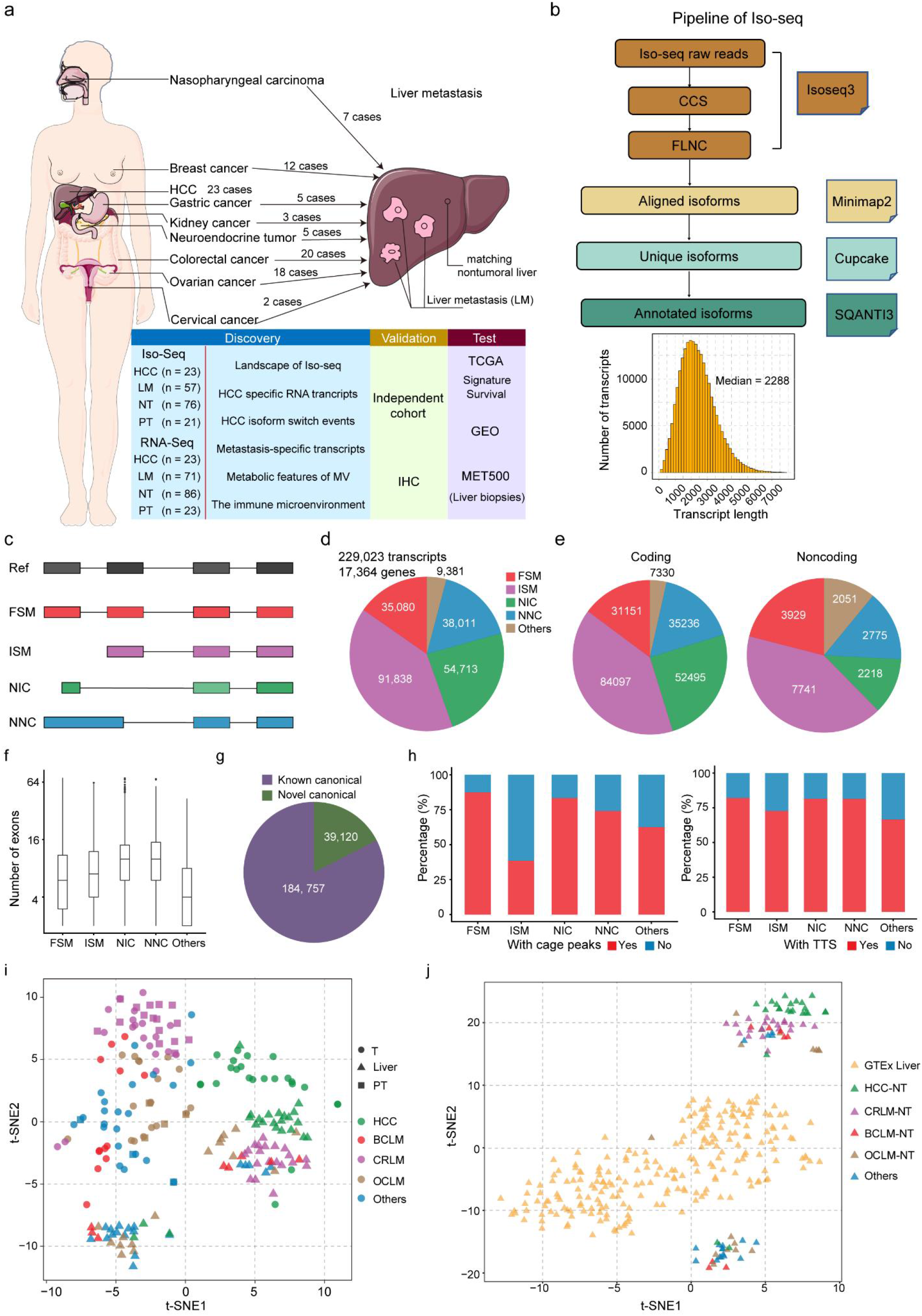
Landscape of long-read transcriptomes in primary and metastatic liver cancers. a. Schematic of primary and metastatic liver cancer isoform profiling by Iso-seq and RNA-seq. b. Isoform calling algorithm for Iso-Seq data. c. Types and illustration of the identified isoforms. (d-e). The percent and number of distinct isoforms in each category from (c) are indicated, including total, coding, and noncoding transcripts. f. Characteristics of novel (NIC and NNC) and known (FSM and ISM) transcripts. NIC and NNC have more exons. g. Proportion annotation and unannotated junctions by comparison with reference transcripts. h. Percent of Iso-seq isoform transcription start sites supported by CAGE (FANTOM5) or transcription termination sites supported by the presence of a poly(A) motif (SQANTI2), plotted per category from (c). i. Global transcript expression patterns of primary and metastatic liver cancers as illustrated by a t-distributed stochastic neighbor embedding (t-SNE) projection. The position of samples within the plot reflects the relative similarity in the expression of transcripts. Samples are color-coded on the basis of their assigned analysis cohort. j. Global transcript expression patterns of the liver samples as illustrated by a t-SNE projection, including HCC-NT, LM-NT and GTEx normal liver samples.

Isoforms obtained with single-molecule real-time circular consensus sequencing were polished using the ToFU (Transcript Isoforms Full length and Unassembled) pipeline (**Fig. 1b**). In total, we identified 229023 unique FL transcript isoforms spanning 17364 annotated genes, with a mean isoform length of 2.3 kb, in liver cancer (**Fig. 1b**). The isoforms were classified into five categories based on their junction match to a reference transcriptome (GENCODE v.35) using SQANTI (**Fig. 1c**). Among the isoforms, 15.3% (35080) were full-splice match (FSM) matching perfectly to known transcripts, 40.1% (91838) were incomplete-splice match (ISM; fewer 5’ exons than reference, but each internal junction agrees), and 23.9% (54713) and 16.6% (38011) were novel in catalogue (NIC; a combination of known splice donors or acceptors that have not been previously cataloged in the same transcript) and novel not in catalog (NNC; at least one splice site not present in reference). The others included isoforms of antisense and intergenic transcripts (**Fig. 1d**). The distributions of transcripts were similar between primary cancer and liver metastases tissues and between HCC and matched non-tumor liver tissues. However, the proportion of each category in the LM-NT group was significantly different from that in the other groups. In addition, the distributions of isoforms were different based on the origin of primary cancers (Additional file 3: Fig. S1a-c). Most of those transcripts were protein-coding transcripts, and the distribution was different between the protein-coding and noncoding transcripts. Of these, 14.8% and 20.99% were FSM, 40.0% and 41.36% were ISM, 25.0% and 11.85% were NIC, and 16.8% and 14.83% were NNC in the protein-coding and noncoding transcripts, respectively (**Fig. 1e**). Compared to known isoforms, novel transcript isoforms (both NIC and NNC) had larger numbers of exons and longer transcript lengths (**Fig. 1f** and Additional file 3: Fig. S1d). Notably, 39120 junctions were not annotated in GENCODE v35 (**Fig. 1g**). We used various quality features provided by SQANTI2 to assess the reliability of the full-length isoforms, including functional genomic evidence such as overlap of 5′ transcript ends with independently published cap analysis of gene expression (CAGE) data and 3′ ends with polyA tails (**Fig. 1h**). Moreover, the number of detected isoforms in samples was highly associated with sequencing reads (Additional file 3: Fig. S1e).

We then compared the distribution of transcripts based on the protein-coding and noncoding transcripts in each group. Overall, novel isoforms accounted for 19% to 29% of the sequenced transcripts in each group (average = 22.9%). Interestingly, similar to the general distribution pattern of total transcripts, the distributions of isoforms were similar regardless of protein-coding and noncoding transcripts, and the proportion of each category in the HM-NT group was significantly different from that in the other groups (Additional file 3: Fig. S1f-o).

To evaluate to what extent tissue-and cancer-specific gene expression was maintained across primary and metastatic lesions, we used the t-distributed stochastic neighborhood embedding (t-SNE) projection to qualitatively visualize isoform expression. It was obvious that tumors were significantly different from non-tumor liver tissues, including non-tumor liver tissues from HCC and metastatic liver cancer patients. In addition, HCC samples were segregated from liver metastasis samples. Compared with HCC, metastatic samples were less well separated based on the type of primary cancer. In addition, most of the primary and matched metastatic samples did not segregate on the basis of the primary cancer type (**Fig. 1i**). Most importantly, the non-tumor liver tissues from HCC patients and LM patients and GTEx normal liver samples were segregated from each other, which indicated that the environment of the healthy liver may be modified for the transformation of HCC or homing of metastatic cells from extrahepatic malignancies (**Fig. 1j**). Taken together, these results indicated that the expression of the isoform showed a tissue-specific transcription pattern and might contribute to the adaptation and colonization of metastatic cancer cells in the liver.

### Profiling of significant differentially expressed RNA transcripts in primary liver cancer

To assess the biological and clinical significance of the isoforms in HCC, we first quantified the expression of isoforms in HCC and HCC-NT, including our Iso-seq HCC samples and previously published nonmetastatic HCC cases (see Methods). Genes were binned into three groups based on our RNA-seq expression levels: low, average, and high based on TPM cut-offs. The average detected total isoforms and novel isoforms (NIC + NNC) increased for genes expressed at low, average and high levels. Novel isoforms were also detected for genes with the lowest expression **(Fig. 2a)**. These data indicate that Iso-Seq detected transcripts even for genes with low expression and that NIC and NNC isoforms from our Iso-Seq data are expressed at appreciable levels. When comparing the gene expression profiles between the tumor and adjacent non-tumor tissues, we identified 10880 differentially expressed transcripts (DETs) after adjusting for multiple testing **(Fig. 2b)**. To further elucidate the significance of gene versus isoform expression patterns, we next examined differentially expressed genes (DEGs) with no isoform expression changes and DETs with no gene expression changes **(Additional file 3: Fig. S2)**. The Venn diagram in **Figure 2c** illustrates the overlap between DETs and DEGs at the gene level. While only 2.4% (119 of 5009) of DEGs had no change in isoform distribution, more than half (55.1%) of DETs belonged to genes with no gene expression changes **(Fig. 2c and Additional file 4: Table S2)**. These DETs, which might have previously been inaccessible and were hidden within comparable gene expression levels, represent an additional dimension in differential expression analysis. Pathway enrichment analysis [Kyoto Encyclopedia of Genes and Genomes (KEGG) and MSigDB Hallmark] for genes with DETs was conducted. Spliced genes with novel isoforms were strongly associated with key HCC pathways, which were overrepresented, such as replication (Myc targets and G2M checkpoint pathway), NF-κB signaling, mTORC1 signaling, and Wnt signaling pathways et al **(Fig. 2d)**.

**Figure 2.**
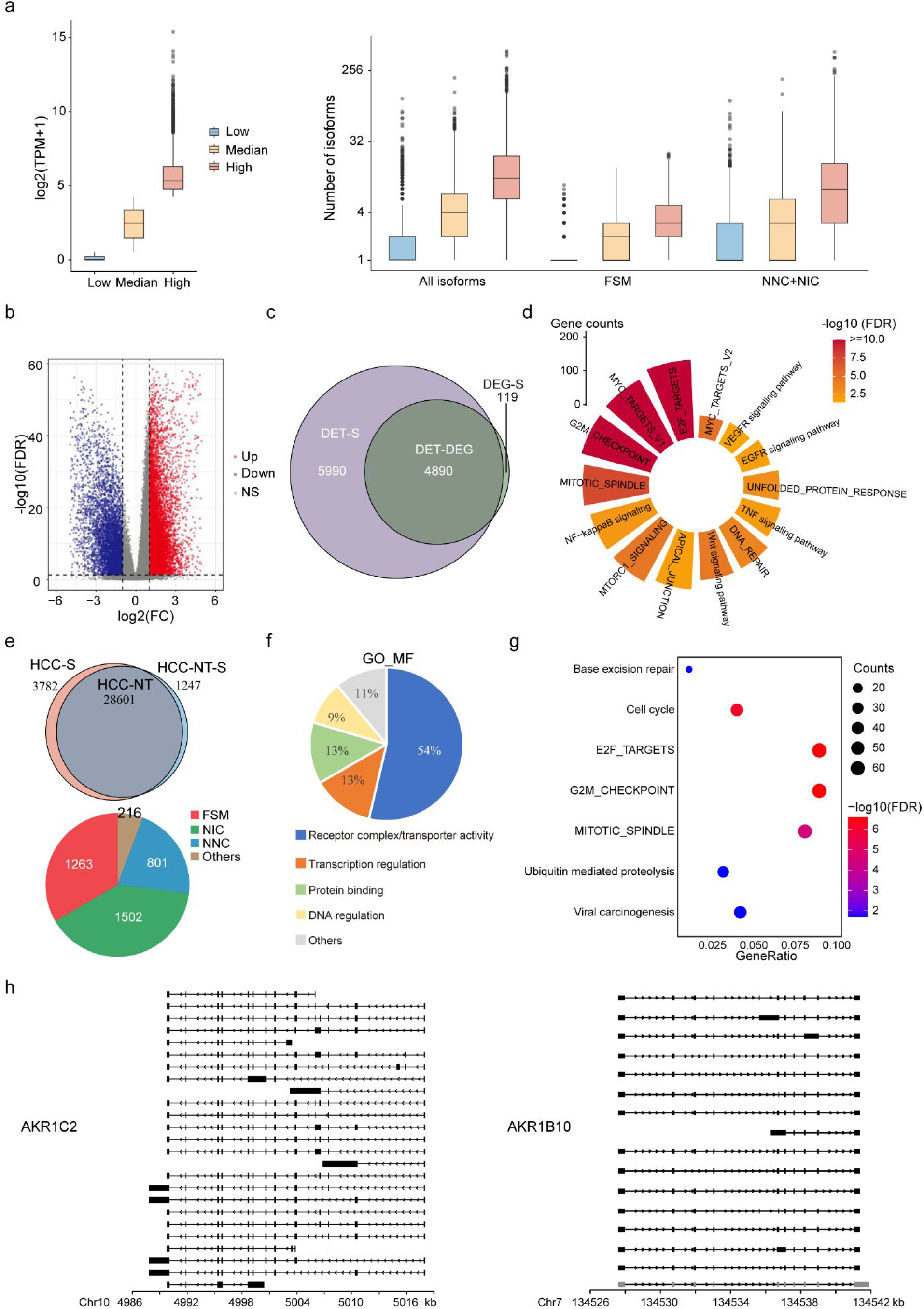
Profiling of significant differentially expressed RNA transcripts between primary HCC and matched non-tumor liver tissues. a. Correlation between gene expression levels from RNA-seq and the number of transcript isoforms detected by Iso-seq. Genes are binned on the basis of quartile expression: low (first quartile), median (second and third quartiles), and high (fourth quartile); b. Volcano plot of DETs in HCC and HCC-NT. Red and blue dots represent transcripts that were significantly upregulated and downregulated (FDR < 0.05), respectively. c. Venn diagram of overlapping DETs and DEGs between HCC and HCC-NT. d. Pathway enrichment analysis for genes with DETs. e. Top, Venn diagram of the number of common and SRTs in HCC and HCC-NT samples. SRTs are defined as a fold change larger than 10 between tumor and paired adjacent non-tumor tissues. Bottom, Distribution of isoform numbers for HCC SRTs. f. GO of HCC SRT-associated genes. g. Pathway enrichment analysis of HCC SRTs. h. Structure of the AKR1C2 and AKR1B10 transcripts in HCC.

Next, we aimed to investigate the specific RNA transcripts (SRTs), that were expressed in HCC but not HCC-NT. We obtained 3782 SRTs in HCC, including 3539 protein-coding RNAs and 243 noncoding RNAs, and most of them were novel transcripts **(Fig. 2e and Additional file 5: Table S3)**. Gene Ontology (GO) analysis of HCC SRTs categorized 80% of enriched molecular function terms as associated with receptor complex/transporter activity, transcription regulation, and protein binding **(Fig. 2f)**. Pathway enrichment analysis revealed that these isoforms were enriched for pathways that are deregulated in HCC, such as the cell cycle, E2F targets, and mitotic spindle **(Fig. 2g)**. The individual oncogenes that had a high gain of novel isoforms in the Iso-seq cancer transcriptome were identified; for example, AKR1C2 and AKR1B10 are often overexpressed in HCC and contribute to hepatocarcinogenesis. In addition to the isoforms in GENCODE v.35, we detected a 2-(AKR1C2) to 2.25-fold (AKR1B10) increase in NIC + NNC isoforms (**Fig. 2h**).

### Clinical significance and expression control of SRTs in primary liver cancer

To assess the clinical relevance of the HCC-SRTs, we applied consensus clustering to classify the patients into two subtypes based on the expression profiles of SRTs. After unsupervised clustering, 50 (13.5%, of 371) HCC patients from the TCGA data set were identified in the SRT-high subtype, whereas the other 321 patients were identified in the SRT-low subtype **(Fig. 3a)**. As expected, the patients from the SRT-high subtype had a significantly worse prognosis and overall survival **(Fig. 3b)**.

**Figure 3.**
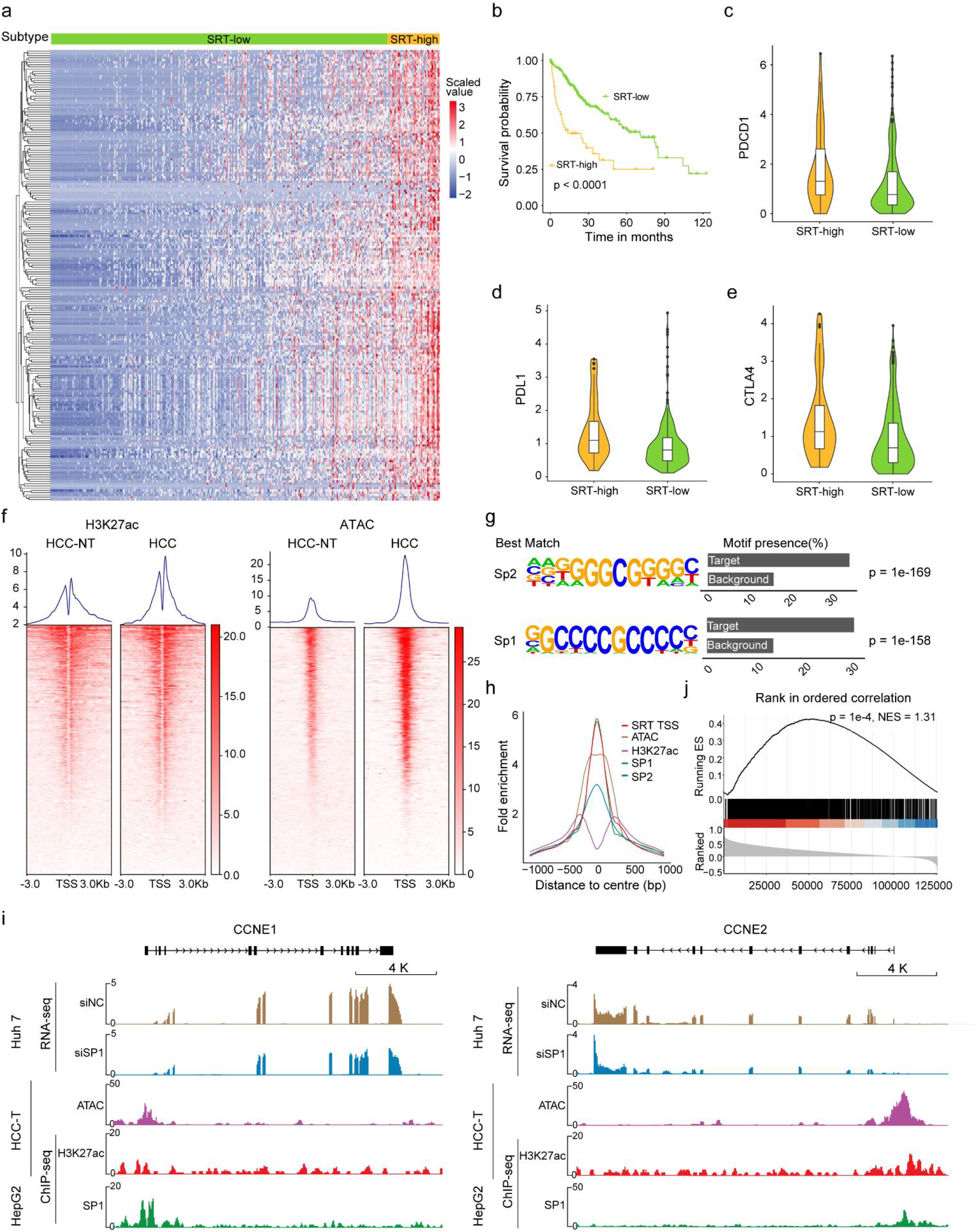
Clinical significance and expression control of the SRTs in primary liver cancer. a. Unsupervised clustering of the HCC SRTs. b. Kaplan-Meier curves show overall survival in SRT-high (red) and SRT-low (green) patients in the TCGA-LIHC cohort. c-e. Differences in the expression of immune checkpoint inhibitors between the SRT-high and -low groups. f. Enrichment of ChIP-seq peaks for H3K27ac and ATAC-seq peaks within 3 kb from the TSSs of HCC SRTs in HCC and HCC_NT. g. Top enriched DNA binding motifs with significant P values identified in a de novo analysis of sequences within 0.5 kb from the TSSs of HCC SRTs. h. Line plots showing ATAC-seq and ChIP-seq signals of H3K27ac, SP1, and SP2 centered at the most enriched motif of SRTs. i. Profiles of SP1 and H3K27ac occupancy and ATAC-seq peaks at the CCNE1 and CCNE2 promoter regions in liver cancer cells and HCC tissues. j. GSEA of HCC SRTs is shown. Transcripts are ranked by the correlation between the expression of SP1 and SRTs. The NES and FDR are shown.

Next, we further attempted to investigate the functional effects of SRT events associated with cell proliferation, EMT, the expression of immune checkpoint inhibitors and the cancer hallmarks of cancer cell stemness, angiogenesis, anti-apoptosis, glycolysis, hypoxia, and inflammation. The results showed that PDCD1 (PD-1), CD274 (PD-L1), and CTLA4 were significantly upregulated in the SRT-high subtype compared to the SRT-low subtype **(Fig. 3c-e)**. The SRT events were positively associated with the majority of cancer hallmarks, such as cell proliferation, stemness, glycolysis, hypoxia, and inflammation **(Additional file 3: Fig. S3a-b)**. However, there were no associations between the SRT events and EMT or angiogenesis. These results suggested that SRTs are associated with poor prognosis and the TME-infiltrating immune response and demonstrated strong functional effects among cancer hallmarks.

To explore the generation of such SRTs, we first used chromatin immunoprecipitation sequencing (ChIP-seq) and assay for transposase-accessible chromatin (ATAC) sequencing to investigate variations in the promoter landscapes between HCC and HCC-NT. Analysis using an antibody against H3K27ac showed that the genome regions around loci encoding SRTs had enriched H3K27ac deposition and more chromatin accessibility in HCC compared to HCC-NT **(Fig. 3f)**. To determine whether some key transcription factors were involved in the generation of SRTs, we conducted a de novo binding motif analysis to examine all of the gained promoters in HCC samples. Interestingly, motifs for the specificity protein (Sp) family, including SP1 and SP2, were highly enriched in the gained promoters in HCC **(Fig. 3g)**. It is well documented that Sp family proteins play a significant role in cancer development and progression[15]. Moreover, SP1 can generate tissue-specific gene expression programs with or without tissue-specific transcription factors. It could be modified to increase binding/transactivation at a specific site, and epigenetic changes could alter the availability of the binding sites[16]. By analyzing the ChIP-seq data for SP1 and SP2 in the liver cancer cell line HepG2, we found that most of the SRTs were bound directly by SP1 and SP2 in their promoters near transcription start sites (TSSs), which have strong H3K27ac occupancy and open chromatin accessibility **(Fig. 3h-i and Additional file 3: Fig. S3c)**. Furthermore, the GSEA results showed that the SRT signature was significantly associated with the expression of SP1 **(Fig. 3j)**. We then performed RNA-seq to elucidate the underlying transcriptional programs affected by SP1. Interestingly, the transcripts that were most significantly downregulated in SP1 knockdown Huh 7 cells compared with control cells were enriched in the SRT signature **(Additional file 3: Fig. S3d)**. Together, these results indicated that epigenetic modification, chromatin accessibility and Sp family proteins play essential roles in the expression control of SRTs in HCC.

### Landscape of isoform switching events in primary liver cancer

As mentioned above, most DETs belonged to genes with no gene expression changes, and we compared the numbers of transcripts identified in our Iso-Seq and in the TSSs of each gene in the database. The results showed that DET-specific genes had a significantly larger number of transcripts than DEGs and had a large number of multiple TSSs **(Fig. 4a-b)**. These data indicate that isoform switching events occurred in these DETs with no gene expression changes. Isoform switches are prominent in cancer and are caused by the difference in the functional potential of the two isoforms. Therefore, we performed isoform switch analysis to characterize switch events by comparing the expression of specific isoforms in HCC and HCC-NT. In total, 1438 genes showed significant isoform switching events **(Additional file 3: Fig. S4a and Additional file 6: Table S4)**; these genes were generally associated with signal transduction and metabolism and were part of known cancer gene signatures **(Fig. 4c-d)**.

**Figure 4.**
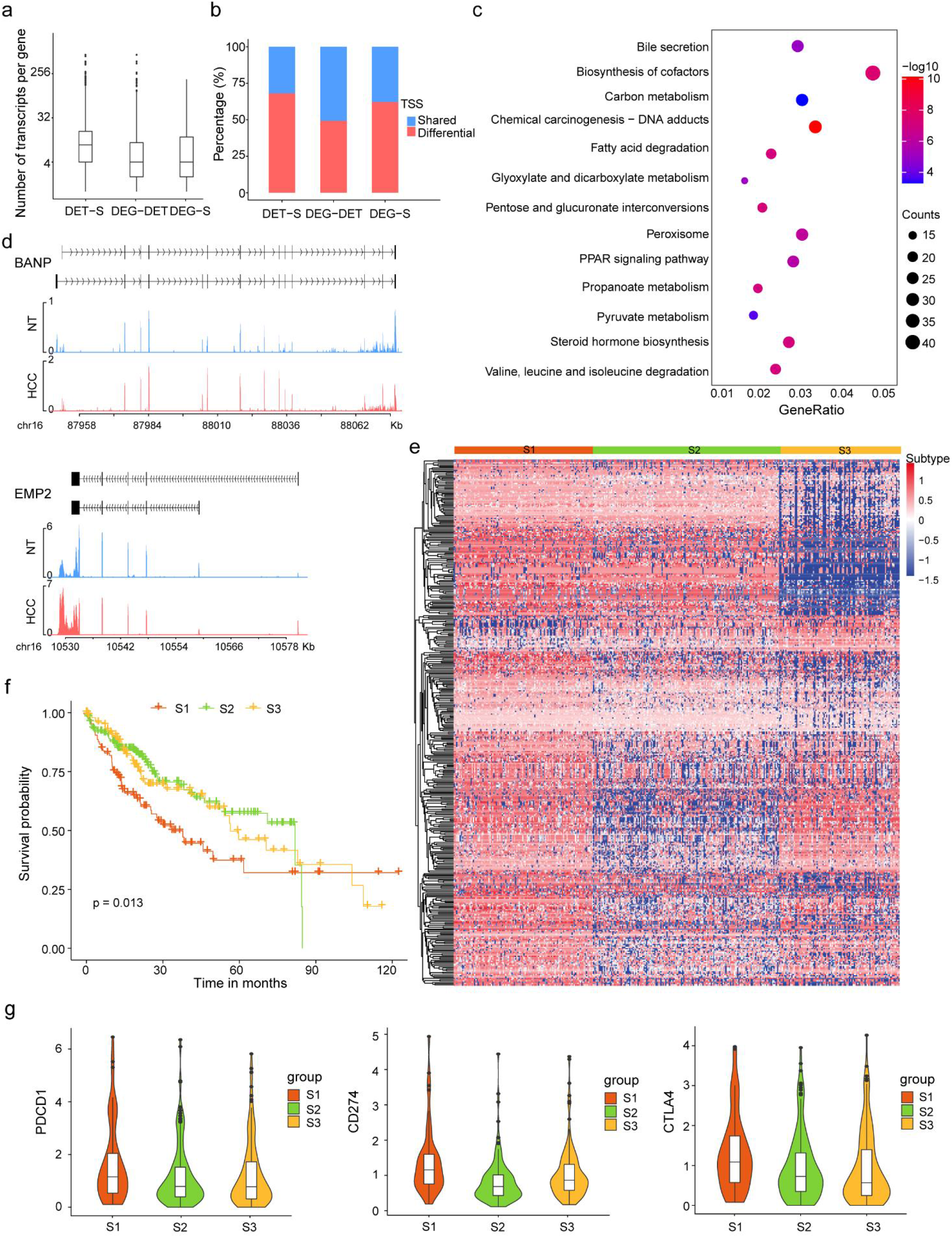
Landscape of isoform switching events in primary liver cancer. a. Comparison of the average number of transcripts per gene for each item. b. Percentage of genes with multiple TSSs for each item. c. Pathway enrichment analysis for isoform switch transcripts. d. Structure of the BANP and EMP2 transcripts in HCC and HCC-NT. e. K-means clustering of the isoform switch ratio. f. Kaplan-Meier curves show overall survival in isoform switch cluster patterns in the TCGA-LIHC cohort. g. Differences in the expression of immune checkpoint inhibitors between types of isoform switch cluster patterns.

We then applied consensus clustering to classify the patients into three subtypes based on the ratio of isoform switching **(Fig. 4e)**. In the prognostic analysis of isoform switch patterns, subtypes revealed a particularly prominent survival disadvantage in Cluster_S1 patients **(Fig. 4f)**. Furthermore, we found that PDCD1 (PD-1), CD274 (PD-L1), and CTLA4 were significantly upregulated in the Cluster_S1 subtype compared to the other subtypes **(Fig. 4g)**. The Cluster_S1 subtype was also positively associated with the majority of cancer hallmarks, such as cell proliferation, stemness, glycolysis, hypoxia, and inflammation **(Additional file 3: Fig. S4b)**. Interestingly, there were no associations between the isoform switch subtypes and EMT or angiogenesis. These results suggested that AS events, including SRTs and isoform switching events, are involved in the early stage of hepatocarcinogenesis.

### Tissue-specific transcript reprogramming is mainly associated with accessible transposable elements and altered TSSs

Interestingly, when evaluating the isoform switch profiles of HCC and matched non-tumor liver tissues in GTEx normal tissues, gene set enrichment analysis (GSEA) showed that the set of significantly downregulated transcripts in HCC was significantly enriched for a liver-specific signature **(Fig. 5a)**. Only the set of liver-specific transcripts was dominantly expressed in matched non-tumor liver tissues, whereas most other tissue-specific transcripts, especially in the testis, were upregulated in HCC, which indicated that such transcript reprogramming to embryonic patterning occurs in HCC initiation **(Fig. 5b and Additional file 3: Fig. S4c)**.

**Figure 5.**
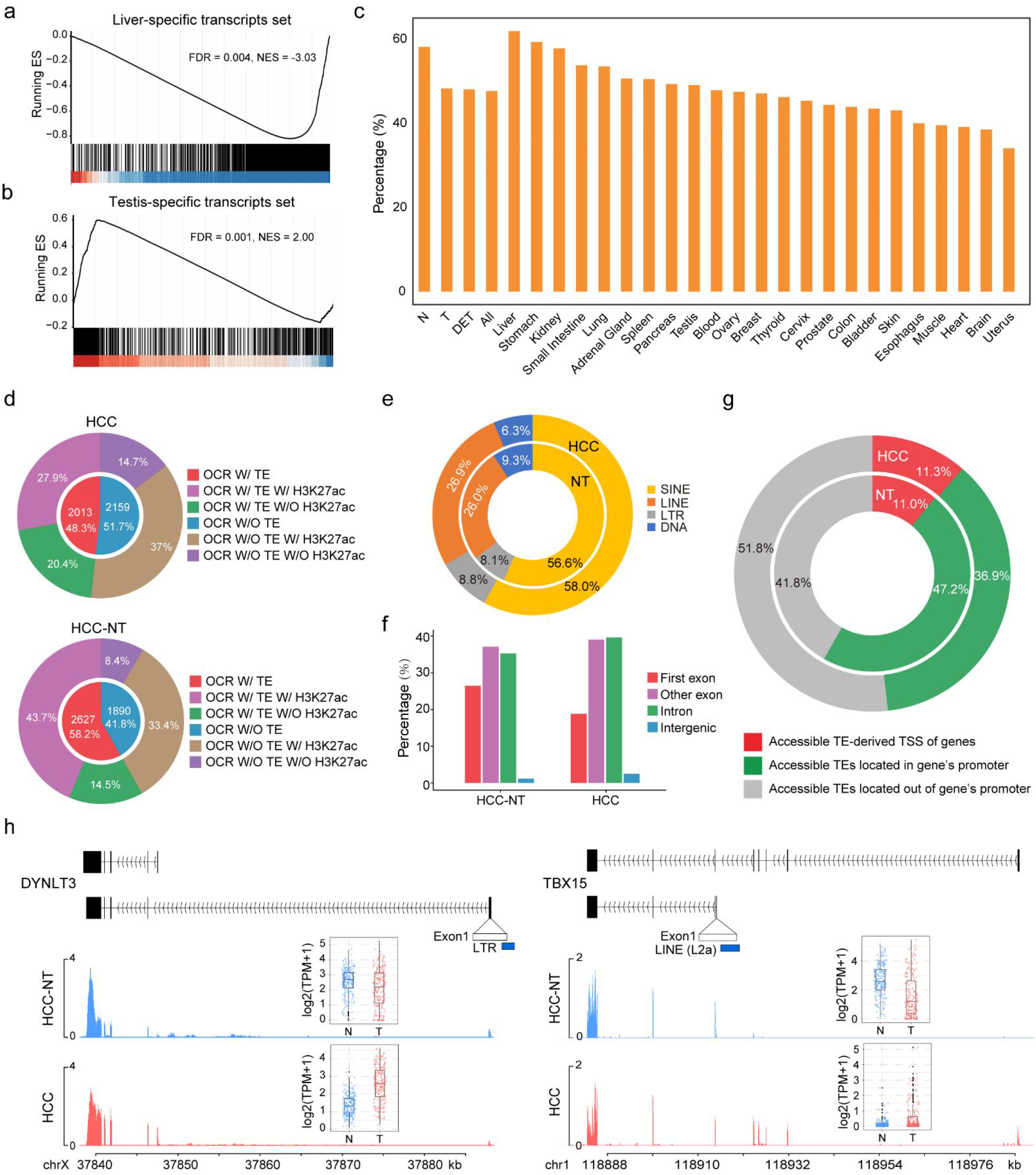
Tissue-specific transcript reprogramming is mainly associated with accessible transposable elements and altered transcription start sites. a-b. GSEA of liver-specific transcripts and testis-specific transcripts as defined from the GTEx database. The transcripts are ranked by the log2-fold change of the TPM values in HCC and HCC-NT samples. c. The percent of isoform switching events containing TEs. d. H3K27ac modification and ATAC-seq peaks identified in HCC and HCC-NT associated with TEs. e. TE class distribution of accessible TEs in HCC and HCC-NT. f. Genomic distribution enrichment of accessible TEs of HCC and HCC-NT in the genome. g. Distribution of accessible TEs in the gene promoter and TE-derived TSS in HCC and HCC-NT. h. Profiles of H3K27ac occupancy around the accessible TE-derived TSSs of isoform switch transcripts in HCC and HCC-NT.

Transposable elements (TEs) are a significant component of eukaryotic genomes and regulate tissue-specific expression during tissue development[17]. We identified approximately 50-60% of isoform switch events from HCC and HCC-NT containing TEs. Interestingly, the tissue-specific transcripts were associated with TEs, and the liver had the most TEs **(Fig. 5c)**. By analyzing ChIP-seq and ATAC-seq data from paired HCC and HCC-NT tissues, 48.3% (HCC) to 58.2% (HCC-NT) of TEs were associated with open chromatin regions (OCRs), and more than 70% of these accessible TEs directly overlapped with the active histone modification H3K27ac **(Fig. 5d)**. On average, more than half of the accessible TEs belonged to the SINE class, and ∼ 26.5% were LINEs. Of the remaining TEs, 8.5% were derived from LTR elements, and less than 8% of the accessible TEs belonged to the DNA class **(Fig. 5e)**. More than half of the accessible TEs overlapped with exons and were specifically located in the first exons, suggesting the potential role of TEs as promoters in tissue-specific transcription reprogramming. Interestingly, we noticed that approximately 40% of accessible TEs were located in intragenic or intergenic regions **(Fig. 5f)**, suggesting that these accessible TEs could be potential enhancers playing a role in isoform switching. Furthermore, we found that approximately half of the accessible TEs were located in the promoter regions (0.5 kb upstream of the TSS) of genes **(Fig. 5g-h)**. In particular, we found that approximately 11% of the accessible TEs directly overlapped with the TSS of genes, suggesting that TEs might have been domesticated as an integral component for RNA Polymerase II recruitment and altered transcription initiation of the downstream gene.

### Long-read transcriptome profiles and characteristics of liver metastases

To explore metastatic transcriptome variations, we analyzed the long-read transcriptomes of primary cancer and liver metastasis samples. We first performed t-SNE to qualitatively visualize the expression of transcripts across our primary cancer and liver metastasis samples, primary cancer samples from TCGA, and normal tissues samples from GTEx. The results showed that the expression of transcripts was highly tissue specific, showing effective segregation of normal and cancer samples among various tissues. Interestingly, these transcripts produced a good separation between primary cancer and metastatic samples, and regardless of the type of primary cancer, metastatic samples clustered together to a certain extent **(Fig. 6a)**. We then evaluated to what extent cancer-specific transcript expression was maintained across metastatic lesions **(Additional file 3: Fig. S5a)**. Compared with normal tissues from GTEx, the DETs in CRLM, BCLM, and OCLM overlapped based on the situation of upregulation and downregulation **(Additional file 3: Fig. S5b-c)**. These overlapping differentially expressed transcript-associated genes were enriched using KEGG and MSigDB cancer hallmarks. Compared with normal original tissues, the transcriptional output was increased for most oncogenic pathways and principal cancer hallmarks. The downregulated transcript-associated genes were enriched in metabolism and cancer-immune responses **(Additional file 3: Fig. S5d-e)**. Next, we compared the DETs between LM and primary cancer tissues to evaluate metastasis-specific transcript variations **(Fig. 6b, Additional file 3: Fig. S5f-g, and Additional file 7: Table S5)**. Metastatic tumors showed a global increase in proliferation, angiogenesis, apoptosis, and hypoxia. Interestingly, spliced genes with novel isoforms were strongly associated with extracellular matrix organization and EMT **(Fig. 6c-d)**. We next examined individual EMT genes that had a high gain of novel splice isoforms in the liver metastasis transcriptome. In total, 23 genes exhibited a twofold increase in NIC + NNC isoforms compared to the reference **(Fig. 6e-f)**. We also compared the DEGs between CRLM, BCLM, and OCLM and previously published nonmetastatic colon cancer (COAD), breast cancer (BRCA), and ovarian cancer (OV) cases from TCGA **(Additional file 3: Fig. S6a)**. The Venn diagram in **Additional file 3: Fig. S6b** shows that the DETs included most DEGs. In addition, we then overlapped the upregulated and downregulated DEGs in CRLM, BCLM, and OCLM. As expected, the shared LM deregulated genes from the isoform expression data included most of the DEGs from the gene expression data. Pathway enrichment analysis indicated that fewer cancer-related pathways were enriched using the DEGs from the gene expression data **(Additional file 3: Fig. S6c-f)**, demonstrating the power of performing differential expression analysis using isoform expression data over gene expression data.

**Figure 6.**
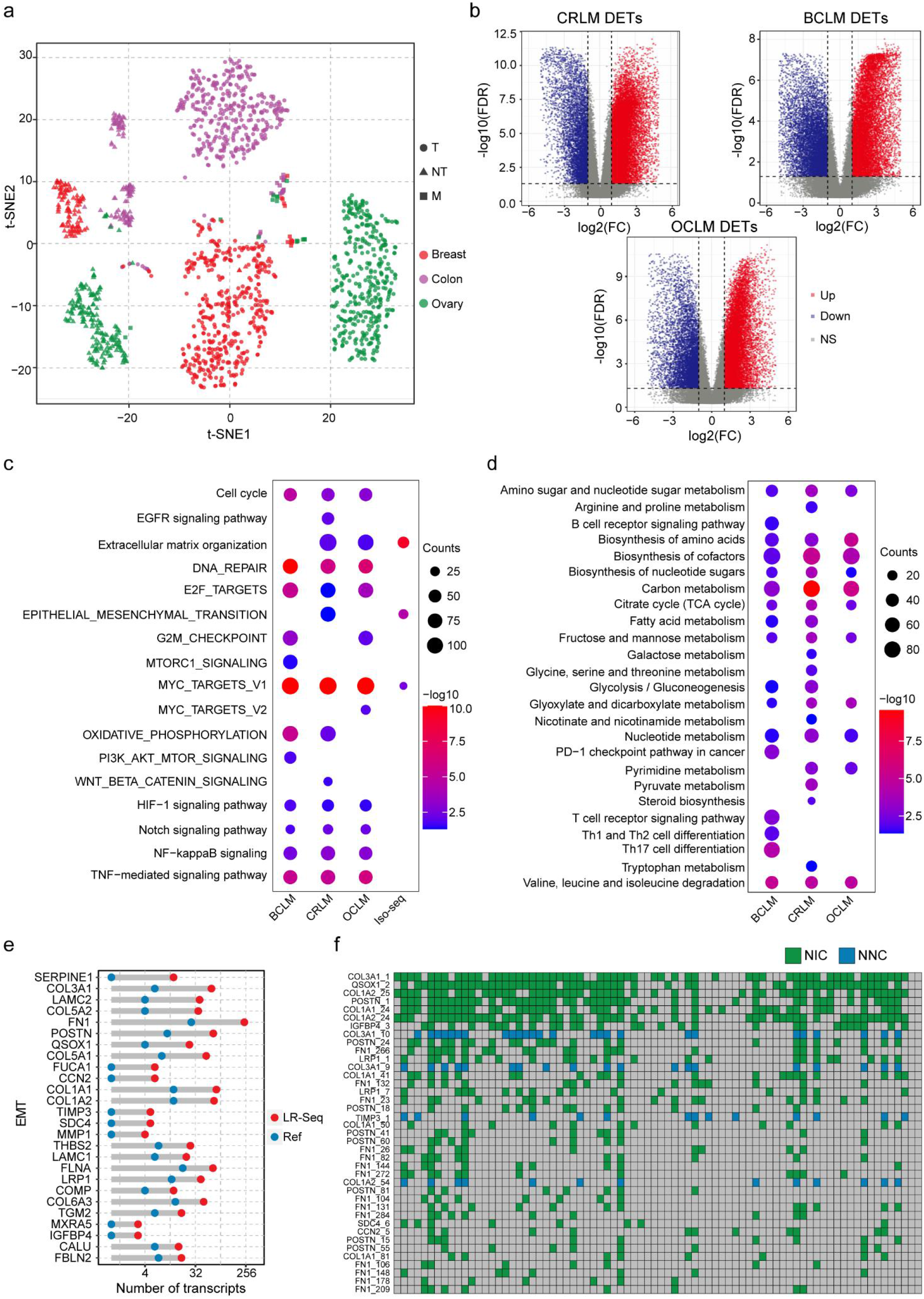
Long-read transcriptome profiles and characteristics of liver metastases. a. Expression patterns of transcripts identified as well differentiated, as illustrated by a t-SNE projection of CRLM, BCLM, OCLM, and primary cancers and the corresponding normal samples. b. Volcano plot of DETs in CRLM, BCLM, and OCLM compared to primary cancer from TCGA. Red and blue dots represent transcripts that were significantly upregulated and downregulated (FDR < 0.05), respectively. c. Pathway enrichment analysis of upregulated transcripts. d. GO analysis of downregulated transcripts. e. Number of novel transcripts compared to annotated GENCODE transcripts for genes of EMT. f. Heatmap shows the expression of novel transcripts of EMT genes in tissues from individual LM patients.

### Metastasis-specific transcripts predict the metastasis and tissue origin of liver metastases

Different tumor types carry specific genetic alterations, and genomic feature analysis could provide precise and pertinent clinical details for disease management. To further understand the effects of metastasis-specific transcripts on the diagnosis of metastasis, we assessed the tumor SRTs in CRLM and BCLM patients. In the context of CRLM, we identified 26 CRLM-SRTs, which were significantly more highly expressed in CRLM patients than in CRC patients from the TCGA data set **(Fig. 7a)**. In addition, these SRTs were barely expressed in other patients from TCGA and in several normal tissues, such as colon, liver, breast, and lung tissues **(Fig. 7b and Additional file 3: Fig. S7a-b)**. To evaluate the diagnostic value of CRLM-SRT expression for liver metastasis of CRC, we further performed receiver operating characteristic (ROC) curve analysis to determine the efficacy of the SRT panel in discriminating liver metastasis in patients with CRC. By random sampling, the combined samples from the CRLM tissue RNA-seq data set and the TCGA colon and rectal cancer RNA-seq data set were divided into a training data set (80%) and a testing data set (20%) for the construction of the CRLM-SRT panel for use in the diagnosis of liver metastasis. The results demonstrated that the CRLM-SRT panel had high accuracy in discriminating CRLM from CRC patients (AUC for training set= 0.935; 95% CI, 0.898 to 0.972; AUC for testing set= 0.966; 95% CI, 0.935 to 0.997) **(Fig. 7c)**.

**Figure 7.**
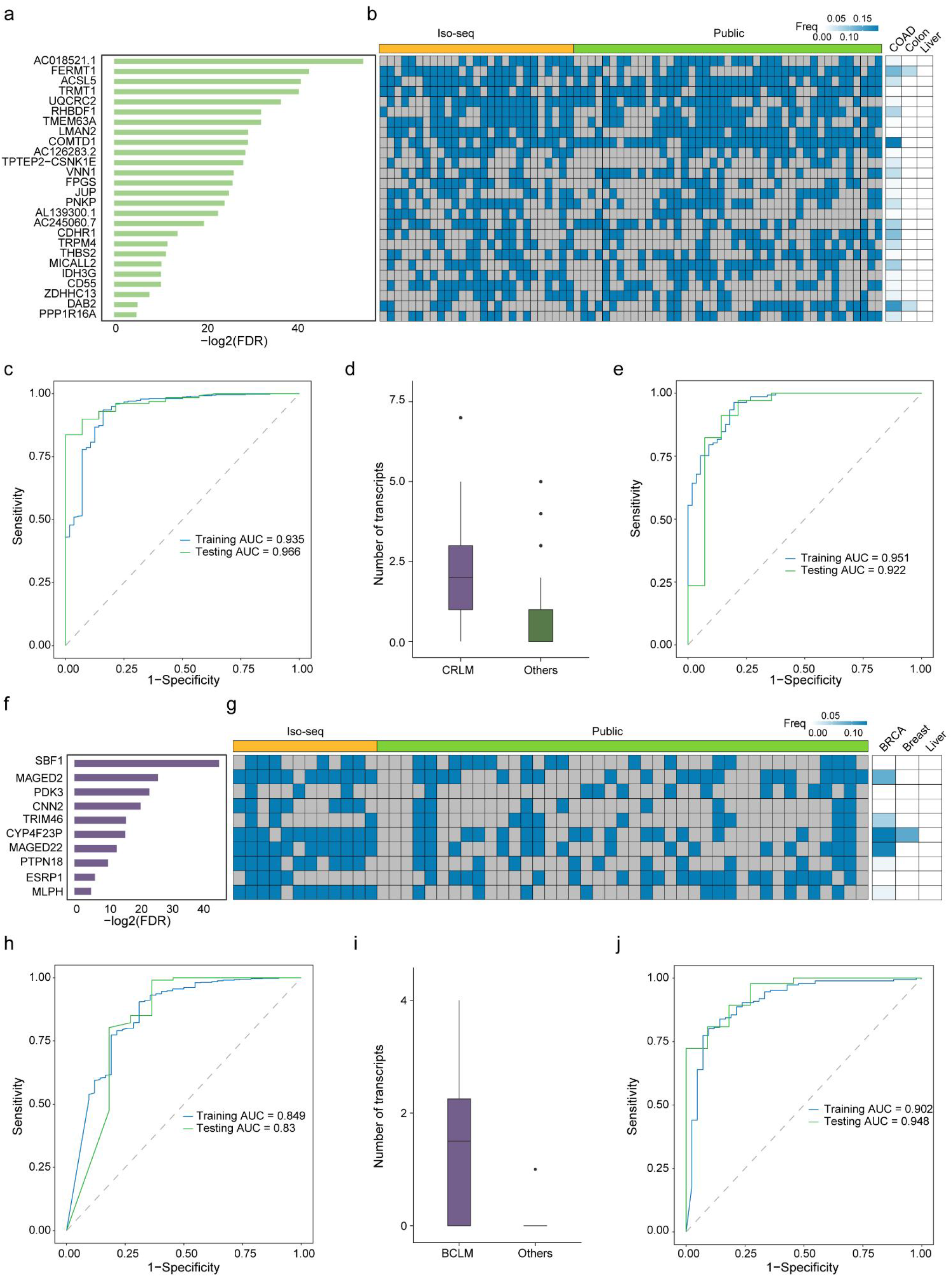
Metastasis-specific transcripts predict the metastasis and tissue origin of liver metastases. a. Barplot shows the statistical significance of 26 CRLM-SRTs. Metastasis-specific transcripts were defined as transcripts with a fold change larger than 10 between metastatic and nonmetastatic tumors. b. Heatmap showing the expression of the indicated novel transcripts in tissues from individual CRLM patients, TCGA-CRC patients, and normal colon and liver tissues. c. Area under the curve (AUC) estimation for the CRLM-SRT panel in the training set and the validation set to discriminate CRLM from CRC. d. Boxplot shows the number of detected SRTs by Iso-Seq in CRLM and LM patients with other tissue origins. e. Area under the curve (AUC) estimation for the CRLM-SRT panel in the training set and the validation set to identify the tumor tissue origin of the colon. f. Barplot shows the statistical significance of 10 BCLM-SRTs. g. Heatmap showing the expression of the indicated novel transcripts in tissues from individual BCLM patients, TCGA-BRCA patients, and normal breast and liver tissues. h. Area under the curve (AUC) estimation for the BCLM-SRT panel in the training set and the validation set to discriminate BCLM from BC. i. Boxplot shows the number of detected SRTs by Iso-Seq in BCLM and LM patients with other tissue origins. j. Area under the curve (AUC) estimation for the BCLM-SRT panel in the training set and the validation set to identify the tumor tissue origin of the breast.

Liver cancer of unknown primary (CUP) is the most common CUP subgroup and has the most dismal prognosis[18]. High expression levels of most CRLM-SRTs were observed in CRLM patients compared with other LM patients **(Fig. 7d)**. We further performed ROC analysis to assess the performance of the CRLM-SRT panel and explore its potential diagnostic utility for LM. The combined samples from our LM tissue RNA-seq data set and the LM tissue RNA-seq from the MET500 data set were divided into a training data set (80%) and a testing data set (20%). For the liver metastatic tumor samples, the overall accuracy of the CRLM-SRT panel was 90.3% to identify the tumor tissue origin of the colon. The AUCs for the CRLM-SRT panel were 0.951 for the training data set and 0.922 for the testing data set **(Fig. 7e)**.

Similarly, we identified 10 BCLM-SRTs that were specifically expressed in BCLM patients compared to BRCA patients from the TCGA data set **(Fig. 7f-g and Additional file 3: Fig. S7c-d)**. The predicted probability was used to construct the ROC curve. The AUCs of the BCLM-SRT panel was 0.849 for the training data set and 0.83 for the testing data set in discriminating BCLM from breast cancer **(Fig. 7h)**. High expression levels of most BCLM-SRTs were also observed in BCLM patients compared with other LM patients **(Fig. 7i)**. Furthermore, the AUCs for the SRT panel were 0.951 for the training data set and 0.922 for the testing data set to predict the tumor tissue origin of the breast in liver metastasis patients **(Fig. 7j)**.

### Altered transcriptome profiles and characteristics of the liver surrounding liver **metastases**

To unravel the implications of the altered liver transcriptomes surrounding liver metastases, we first analyzed the novel transcripts from our Iso-Seq data of the non-tumor livers **(Additional file 8: Table S6)**. We rank-ordered genes based on their ratio of isoform number gain when compared with GENCODE v.35 and selected genes with > twofold increase for pathway enrichment analysis; we found that spliced genes with novel isoforms are strongly associated with metabolism **(Additional file 3: Fig. S8a)**. The individual genes associated with glycine, serine and threonine metabolism that had a high gain of novel splice isoforms were further examined in the liver transcriptomes. In total, 13 genes exhibited a twofold increase in NIC + NNC isoforms compared to the reference **(Fig. 8a)**. Next, we analyzed the transcriptomes between the non-tumor liver tissues surrounding liver metastases and GTEx normal liver tissues to explore the possible factors favoring liver organotropism **(Fig. 8b)**; a total of totally 32595 DETs were identified between these two groups **(Additional file 3: Fig. S8b and Additional file 9: Table S7)**. Pathway enrichment analysis revealed that these DETs were enriched in immunological and metabolic alterations, especially functions associated with the immune response, such as complement and coagulation cascades, TCR signaling, BCR signaling and PD-1 signaling **(Fig. 8c)**. The enrichment of metabolic alteration was associated with the uptake of glucose and central carbon metabolism, including the TCA cycle, glycolysis, hypoxia, and amino acid and fatty acid metabolism. **(Additional file 3: Fig. S8c)**.

**Figure 8.**
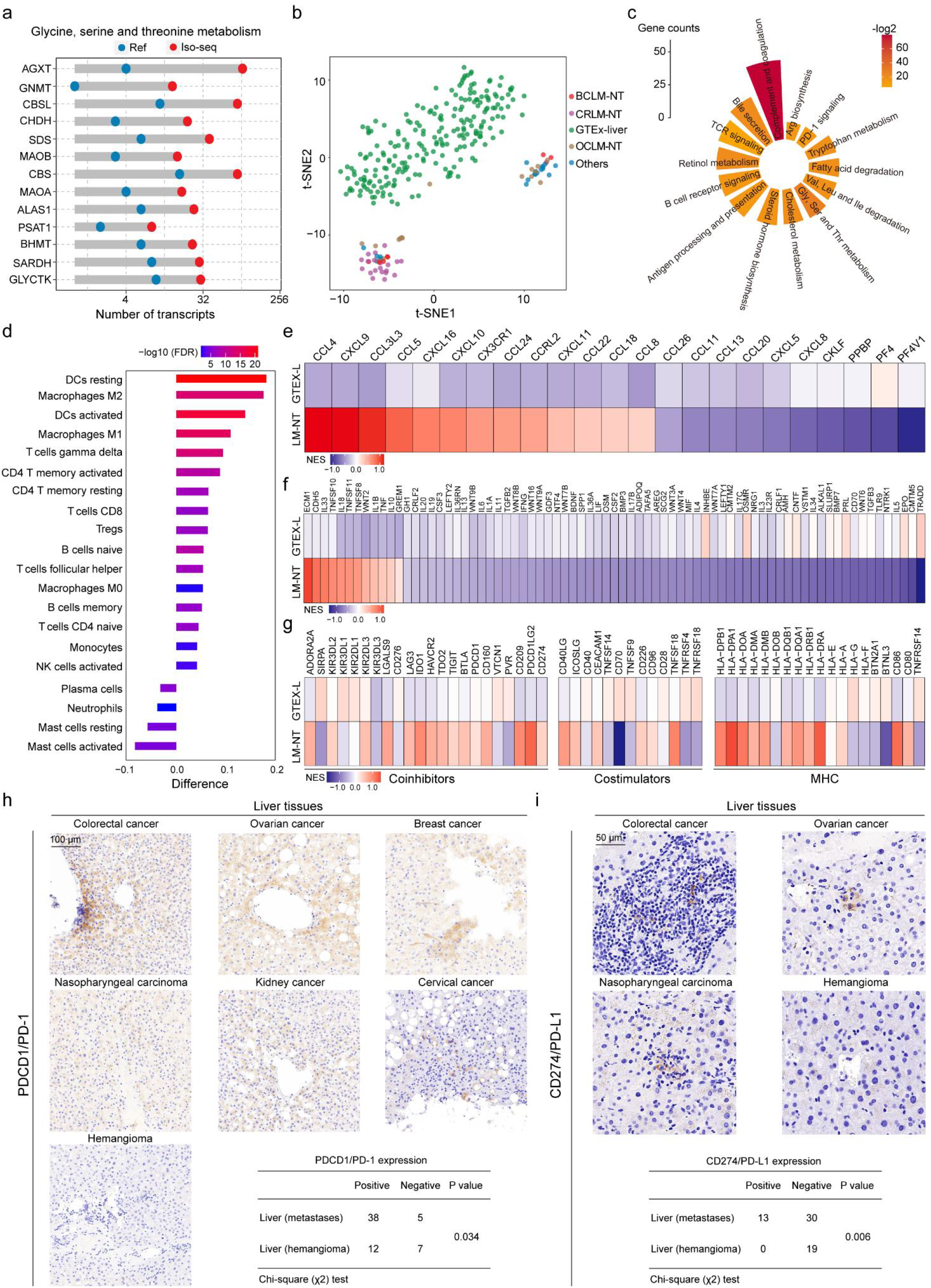
Altered transcriptome profiles and characteristics of the liver in metastatic liver cancer. a. Number of novel transcripts compared to annotated GENCODE transcripts for genes of glycine, serine and threonine metabolism. b. Global transcript expression patterns of the liver samples as illustrated by a t-SNE projection, including LM-NT and GTEx normal liver samples. c. Pathway enrichment analysis of DETs in LM-NT. d. The difference in the relative abundance of infiltrating immune cells between LM-NT and GTEx normal liver samples was calculated by the CIBERSORT algorithm. A difference > 0 indicates that the immune cells were enriched in LM-NT, and the column color represents the statistical significance of the difference. e-f. Change in mRNA expression of chemokines, cytokines, and their receptors, in LM-NT compared to GTEx normal liver samples. g. Change in mRNA expression of MHC molecules, costimulators and coinhibitors in LM-NT compared to GTEx normal liver samples. h-i. Representative IHC images of PD-1 and PD-L1 expression in LM-NT and hemangioma liver tissues. Scale bars, 100 μm or 50 μm.

Furthermore, we analyzed the compositions of different immune cells using a reference microenvironment compendium that included 547 genes representing 22 microenvironment cell subsets in each sample based on the gene expression profile. Altered compositions of immune cell types were observed in the liver tissues surrounding liver metastases. Most cell types, including adaptive immune cells and activated/inactivated innate immune cells, were abundant in the liver surrounding LM, such as resting dendritic cells, M2/M1 macrophages, and T and B cells **(Fig. 8d)**. A significant increase in M2 macrophages and T regulatory cells suggested the role of macrophage activation and immunosuppression in immune escape during liver metastasis. In addition, the liver surrounding LM had higher expression of chemokines, including CCL4 and CCL5, which have been proven to attract monocyte and CD 8+ T cells **(Fig. 8e)**, and highly expressed secreted immunostimulatory and immunoinhibitory cytokines **(Fig. 8f)**. The expression of immune checkpoint molecules after immune stimulation is a potentially important intrinsic immune escape mechanism. Therefore, we referred to a database of costimulatory and coinhibitory molecules to compare these immunomodulators among the clusters. The results demonstrated that the liver surrounding LM had a higher expression of costimulatory molecules (most P < 0.05) and immune checkpoint molecules (most P < 0.05) than the GTEx normal liver **(Fig. 8g)**, suggesting that the liver surrounding LM expressed immune checkpoint molecules to avoid immune killing after immune stimulation. Moreover, we investigated the relationship among immune cells, cytolytic activity (CYT), and the expression of immune checkpoint molecules. The results showed that CYT and the expression of most checkpoint molecules were positively correlated **(Additional file 3: Fig. S8d)**. The deregulated expression of immune checkpoint molecules such as the immune checkpoint inhibitors (ICIs) PDCD1 (PD-1) and CD274 (PD-L1) in adjacent non-tumor liver tissues from LM patients, including CRC, ovarian cancer, breast cancer, nasopharyngeal carcinoma, kidney cancer and cervical cancer liver metastases, was validated by immunohistochemistry (IHC) staining. Consistent with the ICI expression pattern, PD-1-and PD-L1-positive cells showed a significant increase in the liver tissues surrounding LM **(Fig. 8h-i)**. Taken together, these results indicated that the activation of immunosuppressive cells, chemotaxis, and the increase in immunoinhibitory factors might contribute to the extrinsic immune escape of liver metastases.

## Discussion

Long-read sequencing is a potent tool for accurate transcript assemblies and is capable of identifying more novel transcripts. In this study, we defined the full-length isoform-level transcriptome of human primary and metastatic liver cancers and identified isoform-level diversity in HCC and metastasis-associated transcriptome variations in metastatic liver cancers. Moreover, we characterized the specific transcripts to accurately indicate the metastatic potential of primary tumors and the tissue of origin for liver tumors and explored the immunological and metabolic alterations in the liver that make the environment suitable for liver metastases.

A large number of previously undocumented transcripts exhibit critical cellular functions, and their aberrant expression could contribute to carcinogenesis[12–14], which suggests that long-read sequencing technologies provide a better understanding of cancer transcripts. In this study, 70386 isoforms were uncovered in HCC, 32% of which are novel when compared with the reference transcriptome. Several transcripts and transcriptional output were increased for most oncogenic signatures, indicating a global shift towards a cancer-related transcriptional program. Emerging evidence has demonstrated the importance of determining the SRTs of genes in particular physiological and pathological conditions[19, 20]. Our previous studies analyzed normal and cancer RNA-seq samples to identify SRTs that were exclusively expressed in cancer samples and to construct an SRT database across various cancer and tissue types. Herein, we used long-read sequencing to provide more accurate and complete transcriptome-enabling analyses at the SRT resolution and highlight their potential clinical utility in HCC prognosis. Moreover, most of the SRTs contained open chromatin-accessible and H3K27ac-activated promoters, which were occupied by Sp family proteins. The expression of HCC SRTs is positively correlated with Sp1. Sp1, along or cooperating synergistically with tissue-specific transcription factors, can interact at particular regulatory elements, such as promoters and enhancers, to generate tissue-specific gene expression programs, indicating that Sp family proteins might mediate unique promoter changes to activate a set of SRTs in HCC; however, the detailed underlying mechanism needs to be further explored in the future.

Isoform switching events with predicted functional consequences are common in many cancers. Our group and others revealed that some splicing switches contribute essentially to hepatic carcinogenesis and could serve as promising therapeutic targets for HCC[20–22]. However, a comprehensive characterization of the switching events in HCC using long-read sequencing is lacking. Herein, we identified large numbers of events linked to HCC, and patients with specific isoform switch patterns had a worse prognosis. Notably, we discovered reprogrammed tissue-specific transcription, such as the simultaneous loss of liver-specific and gain of other tissue-specific transcription programs, in hepatocarcinogenesis. The intact transcriptional elements of TEs can create novel TSSs to initiate transcription in the host genome[23, 24]. We observed that more than half of the isoform switch events had TEs, and most of them were located in exon regions. Interestingly, we also found that approximately 30-40% of these accessible TEs were located in intron and intergenic regions. These results highlighted the contribution of TEs to tissue-specific transcription reprogramming during carcinogenesis, especially to creating novel tissue-specific transcript expression patterns by acting as alternative TSSs or potential enhancers. Isoform switching in the amino acid sequences among different protein isoforms can have a profound impact on the interaction between the drug and its targets[25, 26]. Overall, the ability to accurately characterize and quantify isoforms of target proteins will undoubtedly provide novel insights for the identification of prognostic markers and inform new potential therapeutic targets for HCC.

Both cancer cell-intrinsic properties (‘‘seed’’) and the congenial ME (‘‘soil’’) are essential for metastasis formation. Progress in the past few decades has greatly enhanced our understanding of the molecular and cellular nature of both the ‘‘seed’’ and ‘‘soil’’[27]. It has long been observed that most cancers show an organ-specific pattern of metastases, and the liver is one of the favored distant metastatic sites for solid tumors[28]. In this study, we investigated metastasis-associated transcriptome variations to explain how metastatic cancer cells with original tissue specificity adapt to the environment of the liver and colonize it. Systematic studies have identified tumor intrinsic factors favoring liver organotropism, including ECM remodeling, cancer stem cells, EMT-MET, and autophagy et. al[29]. We also confirmed that several differentially expressed transcripts-associated genes were enriched in these metastatic oncogenic signatures. Although several spliced isoforms of these genes have been previously identified, current annotations widely used for transcriptome analysis do not contain the level of complexity revealed by our Iso-Seq analysis. A subset of the novel spliced isoforms, leading to novel changes in cellular localization, may play a role in promoting the colonization and metastasis of carcinomas. Targeting these specific transcripts might be counterproductive, and inhibiting signaling pathways could be a promising therapeutic strategy. Furthermore, the current study revealed metastasis-specific RNA transcripts in liver metastatic tumors based on the type of primary cancer, which can predict the metastasis potential in individual patients with CRC and breast cancer. Most importantly, the expression of these specific transcripts achieved a precise prediction of the tissue origin in liver metastases with an overall accuracy of 81.4%, suggesting that the metastasis-specific RNA transcripts can serve as a useful tool for accurately indicating the metastasis potential of primary tumors and clearly identifying the tissue of origin for liver tumors.

Previous studies reported that the tumor microenvironment (TME) of liver metastasis harbors a highly immunosuppressive phenotype and reprogrammed metabolism, induces a systemic loss of antigen-specific T lymphocytes and gains particular benefits in the metabolically active liver[30, 31]. It remains largely unknown how cancer cells modify the environment of the healthy liver for the homing and hosting of metastatic cells. Here, we found that adjacent paracancerous liver tissues are abnormal and represent an immunosuppressive microenvironment that favors the growth of cancer cells. Various circulating factors from cancer cells, such as inflammatory cytokines and chemokines, not only enhance cancer cell proliferation, but also establish the premetastatic niche[32]. M2 macrophages can promote tumor growth and liver metastases[33]. Antigen presentation by immune cells can preferentially lead to immune tolerance via the expression of PD-L1 and engagement of PD-1 on T cells, leading to T-cell exhaustion[34]. This status of immune tolerance further contributes to a pro-metastatic microenvironment[30]. In addition, the metabolic environment can shape the immune response in the liver, enabling tumor cells to escape the mechanisms of immunosurveillance[35, 36]. This study indicated that the activation of the chemokine and cytokine signaling pathways, combined with immune evasion and metabolic reprogramming, plays an important role in the establishment of the premetastatic niche in the liver. How the hepatic microenvironment adapts to the homing and hosting of metastatic cells characterized by these immunological or metabolic alterations to alter immunosurveillance still needs to be further studied, especially using single-cell sequencing to illustrate the immune-metabolic microenvironment of the cancerous liver. Interestingly, liver metastatic patients with non-small cell lung cancer (NSCLC) that had been treated with nivolumab (anti-PD1 antibody) had improved overall survival and progression-free survival[37]. In the present study, we found that the expression of PD-1 and PD-L1 was upregulated in non-tumor liver tissues from a broad range of cancer patients with liver metastasis, which might be the basis for immunotherapy in patients with metastatic liver disease. With more research into the molecular underpinnings of different tumor types, immune checkpoint inhibitors will hopefully continue to be introduced into the clinical setting to treat patients with liver metastases.

## Conclusions

Here we comprehensively surveyed the full-length transcriptome landscapes of primary and metastatic liver cancers at transcript resolution. SRTs are frequently expressed, and isoform switching events often occur in HCC with clinical implications and immunological and metabolic alterations to help cancer cells metastasize to the liver. Our results strongly highlight that full-length transcriptome profiling represents an underexplored research area that may yield novel biological insights and biomarkers. The metastasis-specific transcripts that predict metastatic risk and identify the primary sites of CUP LM patients would help improve the clinical care and outcome of LM patients.

## List of abbreviations

HCC: Hepatocellular carcinoma
CRC: colorectal cancer
EMT: epithelial to mesenchymal transition
AS: alternative isoforms
Iso-Seq: isoform-sequencing
PT: primary tumor
LM: liver metastases
NT: non-tumor
CRLM: colorectal cancer liver metastases
BCLM: breast cancer liver metastases
OCLM: ovarian cancer liver metastases
FSM: full-splice match
ISM: incomplete-splice match
NIC: novel in catalogue
NNC: novel not in catalog
DET: differentially expressed transcript
DEG: differentially expressed gene
SRT: specific RNA transcript
ChIP-seq: chromatin immunoprecipitation sequencing
ATAC: assay for transposase-accessible chromatin
TSS: transcription start site
GSEA: gene set enrichment analysis
TE: Transposable element
COAD: Colon Cancer
BRCA: breast cancer
OV: ovarian cancer
ROC: receiver operating characteristic
CUP: cancer of unknown primary

## Declarations

### Ethics approval and consent to participate

This study was approved by the Ethics Committee of the Fudan University Shanghai Cancer Center (approval number: 2011-ZZK-33). Institutional review board approval was obtained and each patient provided written informed consent. All analyses were performed in accordance with local and international regulations for research ethics in human subject research. This study conformed to the principles of the Helsinki Declaration.

### Consent for publication

Not applicable.

### Availability of data and materials

The raw data of transcriptome and Iso-seq reported in this paper can be accessed from the Genome Sequence Archive (https://ngdc.cncb.ac.cn/gsa/)[38, 39], using the accession number (HRA003557). All data generated or analyzed during this study are included in this published article and its supplementary information files.

## Funding

Our work was supported by grants from the National Natural Science Foundation of China (81930123, 81972247, 82172937 and 82121004).

### Authors’ contributions

XH, ZC, and QS conceived and designed the project. ZC and QS carried out bioinformatics analyses; YL and XL performed the experiments. ZC and LL performed IHC; MX and MS analyzed IHC data; YZ, XW, ZS, MS, LW, and YX provided the cancer samples and clinical information. ZC, QS, and XH wrote the paper with comments from all authors. All authors read and approved the final manuscript.

## Acknowledgements

We would like to thank Berry Genomics (Beijing, China) for the long-read library preparation and sequencing.

