## Supplementary Methods for "Long-read Transcriptome Landscapes of Primary and Metastatic Liver Cancers at Transcript Resolution"

### Cell culture

Huh-7 (ATCC, ATCC Number: RCB1366; RRID: CVCL_0336, 2019) was cultured in Dulbecco's modified Eagle's medium (DMEM) (Invitrogen, Carlsbad, CA, USA) containing 10% fetal bovine serum (FBS) (HyClone, Logan, UT, USA) and antibiotics (penicillin and streptomycin, Invitrogen) at 37 °C in 5% CO2. Cells were assessed for mycoplasma monthly via the qPCR analysis using specific primers for detecting mycoplasma. All these cells were recently authenticated by STR analysis.

### Oligonucleotide transfection

SiRNAs or negative control siRNAs (siNCs) were synthesized by RiboBio (RiboBio Biotechnology, Guangzhou, China). Specific siRNA sequences targeting SP1 are as follows (5'-3'): siSP1-1, UGGGAAUUAUGAACUUUACUACC, siSP1-2, GCCUAAUAUUCAGUAUCAAGUAA. Approximately 1-2 × $10^5$ cells were seeded into six-well plates and grown for one day. At 30-40% confluence, cells were transfected with 5 µl of siRNA or siNC (20 µM) using Lipofectamine RNAiMAX (Invitrogen). After 36 h, cells were harvested for further experiments. For RNA-seq, RNA was extracted after 24 h.

### RNAseq

The RNA samples of clinical tissue specimens and Huh 7 cells were extracted using TRIzol reagent (Invitrogen, CA, USA). Sample QC of input Total RNA samples assessed by measuring the RNA Integrity Number (RIN) and concentration using an Agilent 2100 instrument. The sequencing library was prepared using a VAHTSTM Total RNA-seq Library Prep Kit for Illumina ((Vazyme, Nanjing, China)) and was then subjected to paired-end sequencing on the Illumina HiSeq platform. The paired-end reads were aligned to hg38 using STAR. Bam files obtained from STAR were first converted into bedGraph format using BEDTools and were then converted into BigWig files for visualization with UCSC Genome Browser or IGV. 23 pairs of HCCs and adjacent non-tumorous livers were conducted RNA-seq using a polyA-selected strategy. The RNA-seq of 371 HCC tissues and 50 adjacent non-cancerous samples were obtained from The Cancer Genome Atlas (TCGA; https://portal.gdc.cancer.gov) with official authorization. Raw bam files downloaded from the Genomic Data Commons (GDC) data portal were transformed into fastq files using bedtools (version 2.29.2) bamtofastq. Raw RNA-seq data (fastq files) of 200 HCCs and matched non-tumorous livers were downloaded from Gene Expression Omnibus (GEO) under accession numbers GSE77314, GSE94660, GSE124535 and GSE138485. Transcript quantification (transcripts per kilobase million, TPM) of RNA-seq were performed by Salmon (version 1.5.2) with the mapping-based mode.

**PacBio library and single-molecule sequencing**

cDNA was generated from 2 μg of total RNA per sample using the Clontech SMARTer PCR cDNA Synthesis Kit (catalog# 634925 or 634926) according to PacBio's Iso-Seq Template Preparation for the Sequel System. SMRTbell libraries were constructed using the SMRTbell$^{TM}$ Template Prep Kit (Pacific Biosciences, Part No. 100-259-100), then sequenced on the PacBio Sequel II System (BerryGenomics, Beijing, China).

**PacBio data analysis**

Sequence data were processed using the SMRT Analysis software (IsoSeq 3) in PacBio SMRT Analysis v6.0 to obtain high-quality, full-length transcript sequences, followed by downstream analysis. Sequence was processed by using Iso-Seq (version 3.4.0) workflow to generate full-length reads. Briefly, raw data of sub-reads were merged to circular consensus sequence (CCS) reads with minimum predicted accuracy in 0.9. Full length reads were generated through removal of 5' and 3' cDNA primers by using lima with default parameters. Artificial concatemers reads and polyA tails were then removed by using isoseq3 refine to generate full-length, non-chimeric

(FLNC) reads. FLNC reads were then clustered into high-quality transcripts by using isoseq3 cluster. Filtered transcripts were mapped to the human genome (hg38, GENCODE v35) using minimap2 (version 2.17). Isoforms were subsequently collapsed and correlated using cDNA_Cupcake (version 19.0.0) and SQANTI3 (version 1.6). Transcripts in all samples were merged into a single non-redundant gtf file using gffcompare (version 0.11.2) and annotated using gffcompare and SQANTI3.Next, we filtered the transcripts without junctions support from RNA-Seq. Finally, transcripts detected in at least two biological replicates by Iso-Seq were subsequently used for gtf reference in RNA-seq quantification.

**Differential expression analysis and pathway enrichment analysis**

We utilized the Wilcoxon rank-sum test to assess the statistical difference between tumor and paired normal samples and defined significantly differential expressed isoforms as $|\log2(FC)| > 1$ and $FDR < 0.05$. The functional enrichment and gene set variation analyses were performed by the R package “clusterProfiler” (version 3.16.1) (22455463) as previous of described (35286894).

**Isoform-based clustering**

We performed k-means clustering by the R package “ConsensusClusterPlus” (version 1.52.0). Empirical cumulative distribution CDF plots were generated to determine the optimal number of isoform-based HCC subtypes.

**Machine learning based on random forest**

The harmonized CRLM and CRC patients (n = 713) were randomly grouped into training (n = 570) and held-out testing (n = 143) sets in a 4:1 ratio. Both the training set and the test had the same cancer type distribution. Selected 26 SRTs were employed to build the random forest model by using the R package randomForest (version 4.6.14) with 5-fold cross validation in training set. Prediction of the test cohort was performed with the same parameters. Performance of machine learning model based on SRTs was assessed by receiver operating characteristic (ROC) curve. For BCLM and BRCA cohorts, harmonized patients (n = 1093) were randomly classified into training (n = 874) and held-out testing (n = 219) sets. Ten SRTs were used to construct the random forest model. Models to determine tumor tissue of origin (TOO) were processed in the same procedure.

**Calculation of microenvironment cell abundance**

The gene signatures of LM22 (22 types of immune cells) were obtained from CIBERSORT. Single-sample gene set enrichment analysis (ssGSEA) [28] was performed to calculate the relative level of each immune cell by the transcriptome of samples.

**ChIP-seq library preparation and sequencing**

ChIP-seq for HCC and matched non-tumor liver tissues were performed followed the previously published protocol [1]. Briefly, fresh liver cancer or normal tissues were transferred to a clean 1.5-mL tube containing 250 μL of ice-cold PBS and homogenized to yield chunks 0.5 $mm^3$. The tissues were cross-linked for 15 min with 1% formaldehyde, followed by the addition of 0.15M glycine to terminate crosslinking. Then washed and gathered the tissues into new tubes. Chromatin was digested into DNA fragments using 0.5 μL Micrococcal Nuclease. The complex of protein and DNA was extracted and resuspended in SimpleChIP Chromatin IP buffers (Cell Signaling Technology, Danvers, Massachusetts, USA), and was incubated with magnetic beads (ThermoFisher Scientific, Shanghai, China) conjugated with 1 μg of the antibody against H3K27ac (Active motif, Shanghai, China, RRID: AB_2561016). The target DNA were washed and eluted using MinElute Spin Columns (Qiagen, Hilden, Germany) for DNA sequencing.

ChIP-sequencing libraries were prepared by using the NEBNext Ultra DNA Library Prep Kit for Illumina (NEB Cat. No. E7370) according to the manufacturer's instructions. After barcoding, pooled DNA was sequenced (HiSeq 1500, Illumina) to achieve a minimum of $2 \times 10^7$ aligned reads per sample. For ChIP-seq analyses, 150-bp paired-end reads were aligned to the reference human genome using Bowtie with standard alignment parameters. PCR duplicates were marked with the Picard "Mark Duplicates" utility and removed from further analysis. Bam files were converted to BigWig files using deepTools. The peak distribution along genomic regions of genes of interest were visualized with IGV.

**ATAC-seq**

Fresh liver cancer or non-tumor liver tissues were cut into 1-2 mm pieces and digested using collagenase. Grinding and counting to isolate 50000 living cells. Transpose, purify and PCR steps were performed as previously described [2]. All deep sequencing was performed on the Illumina HiSeq Xten-PE150 or Illumina HiSeq 2500 platform provided by GENEWIZ (GENEWIZ Suzhou, China). The following tools and versions were used for ATAC-seq data analysis: Trimmomatic, SAMtools, Picard, and Bowtie2. First, Nextera adapter sequences were trimmed from the reads by using Trimmomatic. These reads were aligned to a reference genome using Bowtie2 with standard parameters. Picard was then used to remove duplicate reads. These deduplicated reads were then filtered to retain high-quality (MAPQ $\geq$ 30), non-mitochondrial chromosome, non-Y chromosome, and properly paired (SAMtools flag $0 \times 2$) reads.

**Human Tissue microarray and immunohistochemistry**

Core samples were obtained from representative regions of each tumor based on H&E staining. Duplicate 1-mm cores were taken from different areas of the same tissue block for each case (intratumoral tissue and peritumoral tissue). Serial sections (4-μm-thick) were placed on slides coated with 3-aminopropyltriethoxysilane. The immunohistochemistry analysis was carried out as described previously. The primary mAbs used were rabbit anti-human PD-1 (1:200, Cell Signaling Technology, Danvers, MA, USA, RRID: AB_2798734) and rabbit anti-human PD-L1 (1:100, Cell Signaling

Technology, Danvers, MA, USA, RRID: AB_2728833). The immunostaining intensities was scored semi-quantitatively. All samples were anonymously and independently scored by two investigators. In cases of disagreement, the slides were re-examined, and a consensus was reached by the observers.
