## Supplementary Figures for "Long-read Transcriptome Landscapes of Primary and Metastatic Liver Cancers at Transcript Resolution"

### Fig. S1

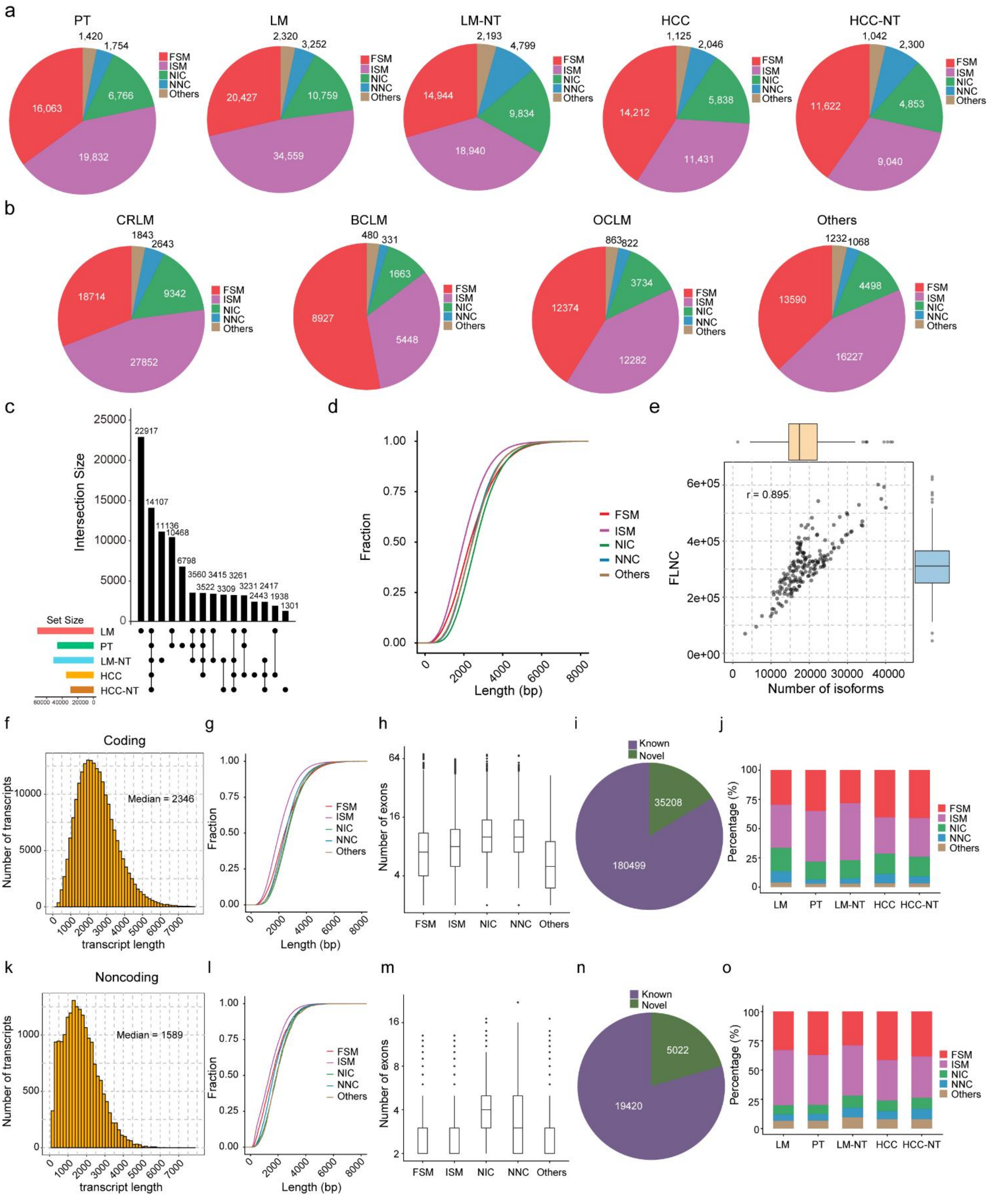


**Fig. S1 Landscape of long-read transcriptomes in primary and metastatic liver**

**cancers**

a. The percent and number of distinct isoforms in each category are indicated in primary and liver metastatic tissues and matched non-tumor tissues. b. The percent and number of distinct isoforms in each category are indicated in different type of liver metastatic tissues. c. Upset plot of shows isoform interactions among five groups of samples. d. Characteristics of novel and known transcripts. e. Correlation between number of detected isoforms and sequencing FLNC reads. f-h. Characteristics of novel (NIC and NNC) and known (FSM and ISM) coding transcripts. i. Proportion annotation and unannotated junctions in coding isoforms. j. The percent and number of distinct coding transcripts in each category are indicated in primary and liver metastatic tissues and matched non-tumor tissues. k-m. Characteristics of novel (NIC and NNC) and known (FSM and ISM) noncoding transcripts. n. Proportion annotation and unannotated junctions in noncoding isoforms. o. The percent and number of distinct noncoding transcripts in each category are indicated in primary and liver metastatic tissues and matched non-tumor tissues.

**Fig. S2**

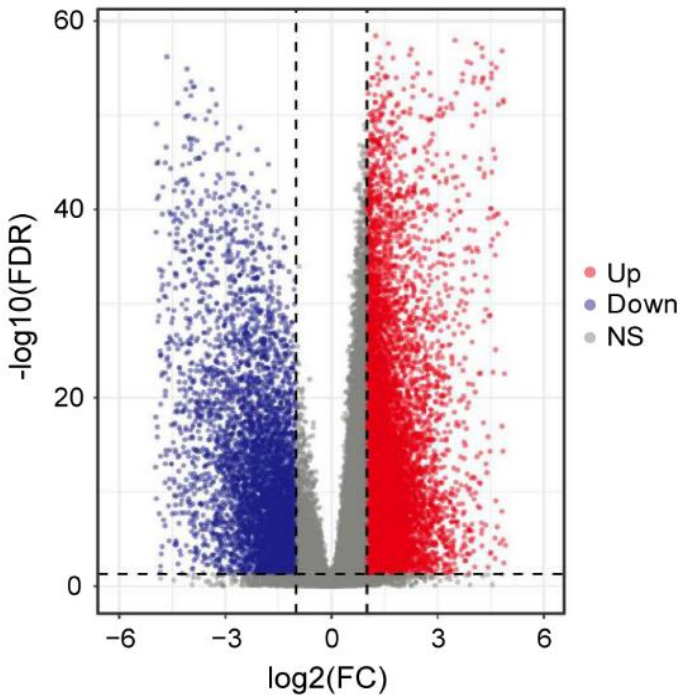


**Fig. S2 Profiling of significant differentially expressed RNA transcripts between primary HCC and matched non-tumor liver tissues**

Volcano plot of DEGs in HCC and HCC-NT. Red and blue dots represent transcripts that were significantly upregulated and downregulated (FDR < 0.05), respectively.

**Fig. S3**

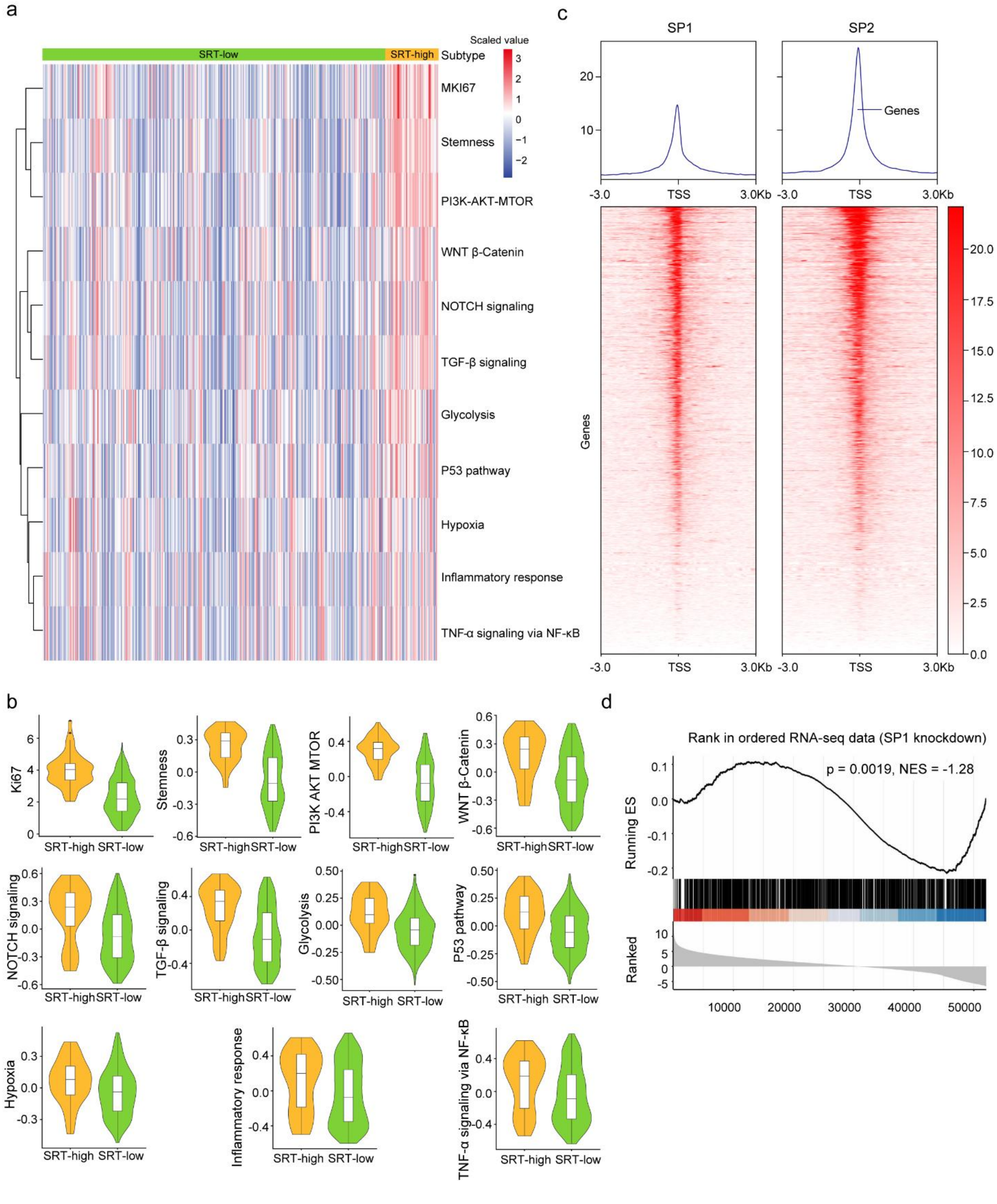


**Fig. S3 Clinical significance and expression control of the SRTs in primary liver cancer**

a. Heatmap visualizing the GSVA enrichment analysis shows the activation states of biological pathways in SRT-high group. b. Differences in the activities of cancer hallmarks between the SRT-high and -low groups. c. Enrichment of ChIP-seq peaks for SP1 and SP2 within 3 kb from the TSSs of HCC SRTs in HCC. d. GSEA of HCC SRTs is shown. Transcripts are ranked by the log2-fold change of the TPM values in Huh 7 siNC and siSP1 cells. The NES and FDR are shown.

**Fig. S4**

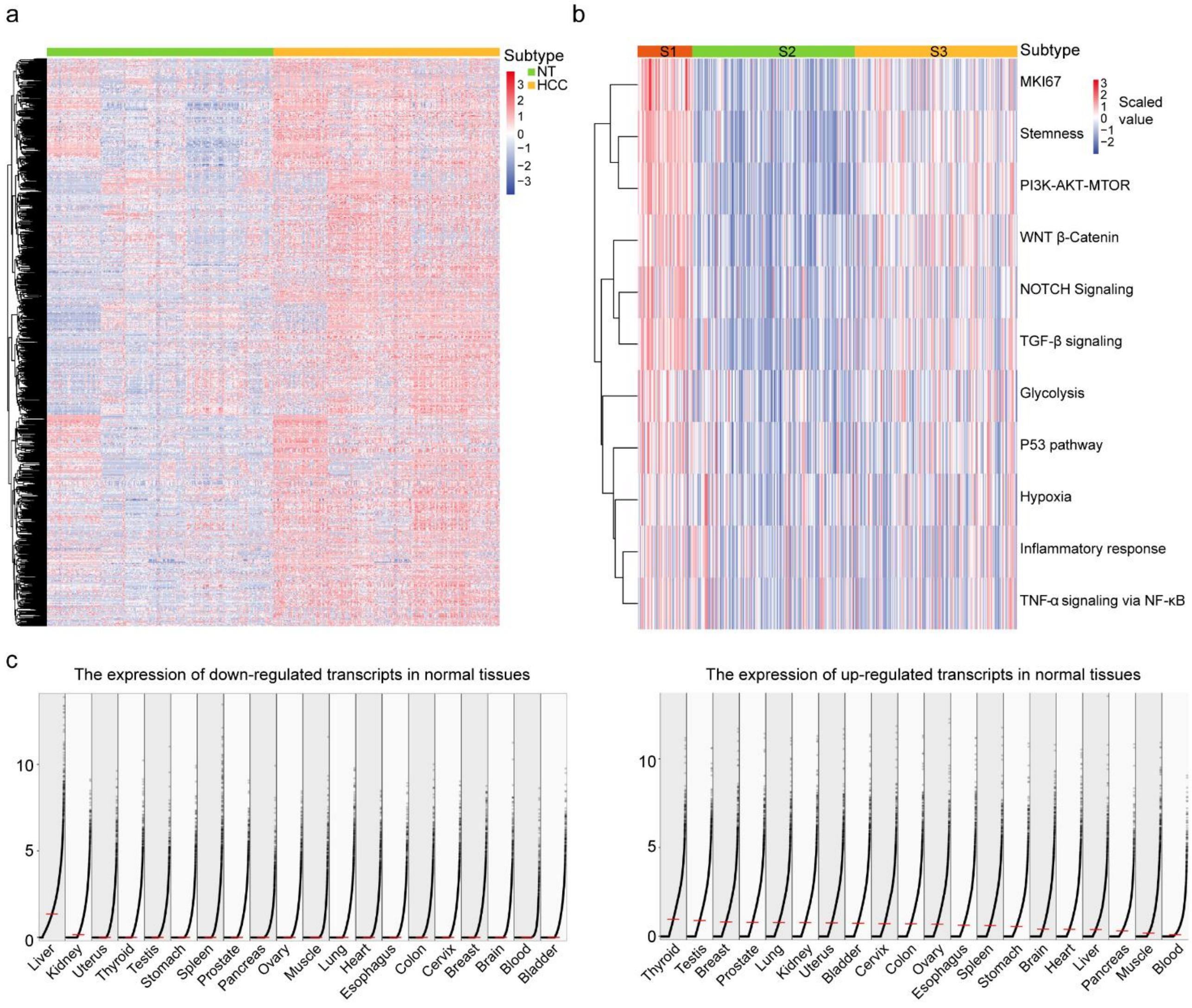


**Fig. S4. Landscape of isoform switching events in primary liver cancer**

a. Heatmap shows the expression levels of isoform switching transcripts in HCC and HCC-NT. b. Heatmap visualizing the GSVA enrichment analysis shows the activation states of biological pathways in isoform switch cluster patterns. c-d. The expression level of isoform switch transcripts among various normal tissues.

### Fig. S5

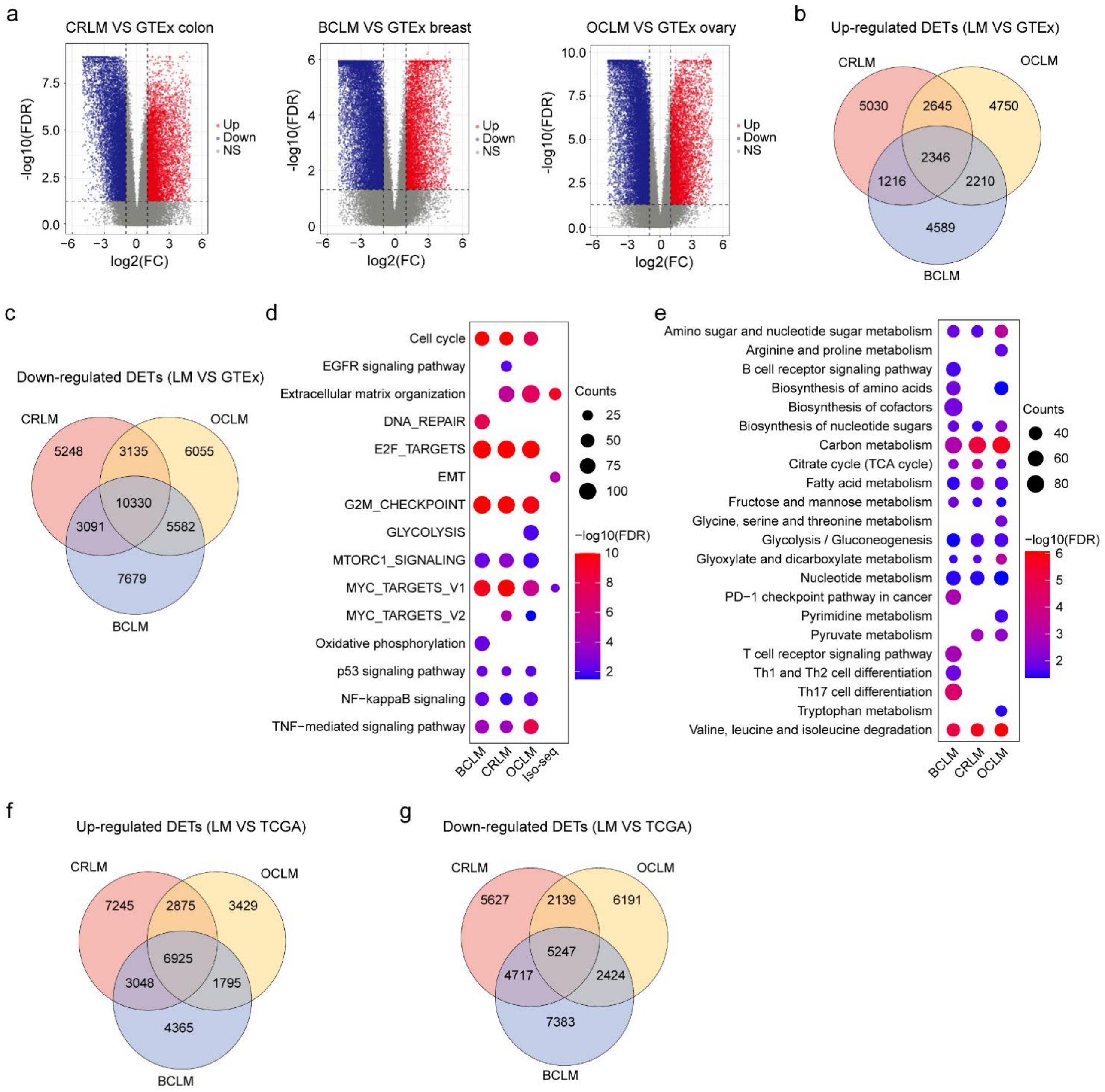


**Fig. S5 DETs in CRLM, BCLM, and OCLM compared to the corresponding normal tissues from GTEx**

a. Volcano plot of DETs in CRLM, BCLM, and OCLM compared to the corresponding normal tissues from GTEx. Red and blue dots represent transcripts that were significantly upregulated and downregulated (FDR < 0.05), respectively. b-c. Venn diagram of overlapping upregulated and downregulated DETs among CRLM,

BCLM, and OCLM. d. Pathway enrichment analysis of upregulated transcripts. e. GO analysis of downregulated transcripts. f. Venn diagram of overlapping upregulated DETs among CRLM, BCLM, and OCLM. g. Venn diagram of overlapping downregulated DETs among CRLM, BCLM, and OCLM.

### Fig. S6

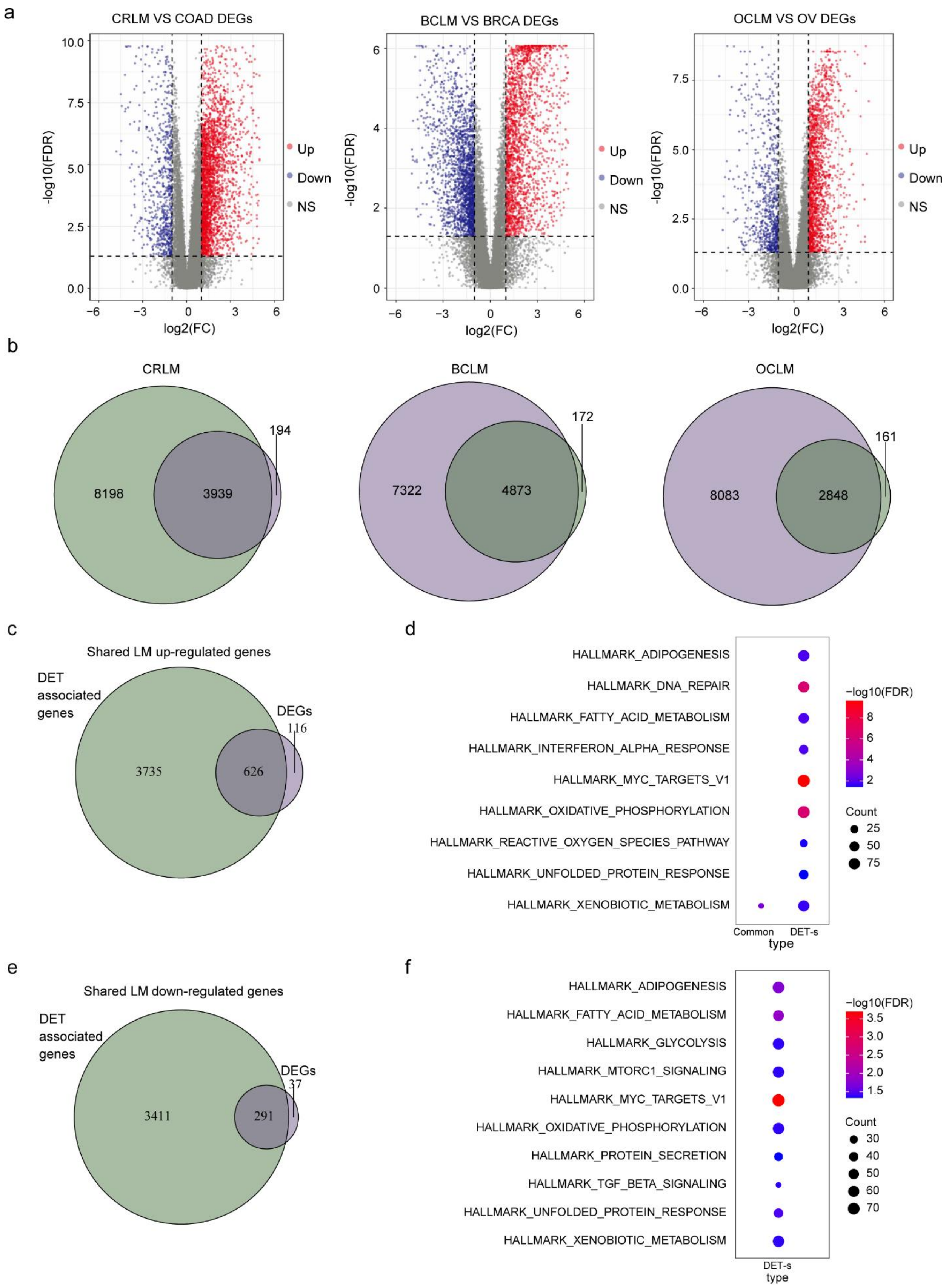
a
CRLM VS COAD DEGs
BCLM VS BRCA DEGs
OCLM VS OV DEGs
-log10(FDR)
log2(FC)
Up
Down
NS
b
CRLM
8198
3939
194
BCLM
7322
4873
172
OCLM
8083
2848
161
c
Shared LM up-regulated genes
DET associated genes
DEGs
3735
626
116
d
HALLMARK_ADIPOGENESIS
HALLMARK_DNA_REPAIR
HALLMARK_FATTY_ACID_METABOLISM
HALLMARK_INTERFERON_ALPHA_RESPONSE
HALLMARK_MYC_TARGETS_V1
HALLMARK_OXIDATIVE_PHOSPHORYLATION
HALLMARK_REACTIVE_OXYGEN_SPECIES_PATHWAY
HALLMARK_UNFOLDED_PROTEIN_RESPONSE
HALLMARK_XENOBIOTIC_METABOLISM
Common
DET-s
type
−log10(FDR)
Count
e
Shared LM down-regulated genes
DET associated genes
DEGs
3411
291
37
f
HALLMARK_ADIPOGENESIS
HALLMARK_FATTY_ACID_METABOLISM
HALLMARK_GLYCOLYSIS
HALLMARK_MTORC1_SIGNALING
HALLMARK_MYC_TARGETS_V1
HALLMARK_OXIDATIVE_PHOSPHORYLATION
HALLMARK_PROTEIN_SECRETION
HALLMARK_TGF_BETA_SIGNALING
HALLMARK_UNFOLDED_PROTEIN_RESPONSE
HALLMARK_XENOBIOTIC_METABOLISM
DET-s
type
−log10(FDR)
Count

**Fig. S6 DETs and DEGs in CRLM, BCLM, and OCLM compared to the corresponding primary cancer from TCGA data set**

a. Volcano plot of DEGs in CRLM, BCLM, and OCLM compared to the primary cancer from TCGA. Red and blue dots represent transcripts that were significantly up- and down-regulated (FDR < 0.05), respectively. b. Venn diagram of overlapping DETs and DEGs in CRLM, BCLM, and OCLM, respectively. c. Venn diagram of overlapping shared LM up-regulated genes (CRLM, BCLM, and OCLM) from DETs and DEGs. d. Pathway enrichment analysis of shared LM upregulated genes. e. Venn diagram of overlapping shared LM down-regulated genes (CRLM, BCLM, and OCLM) from DETs and DEGs. f. Pathway enrichment analysis of shared LM down-regulated genes.

### Fig. S7

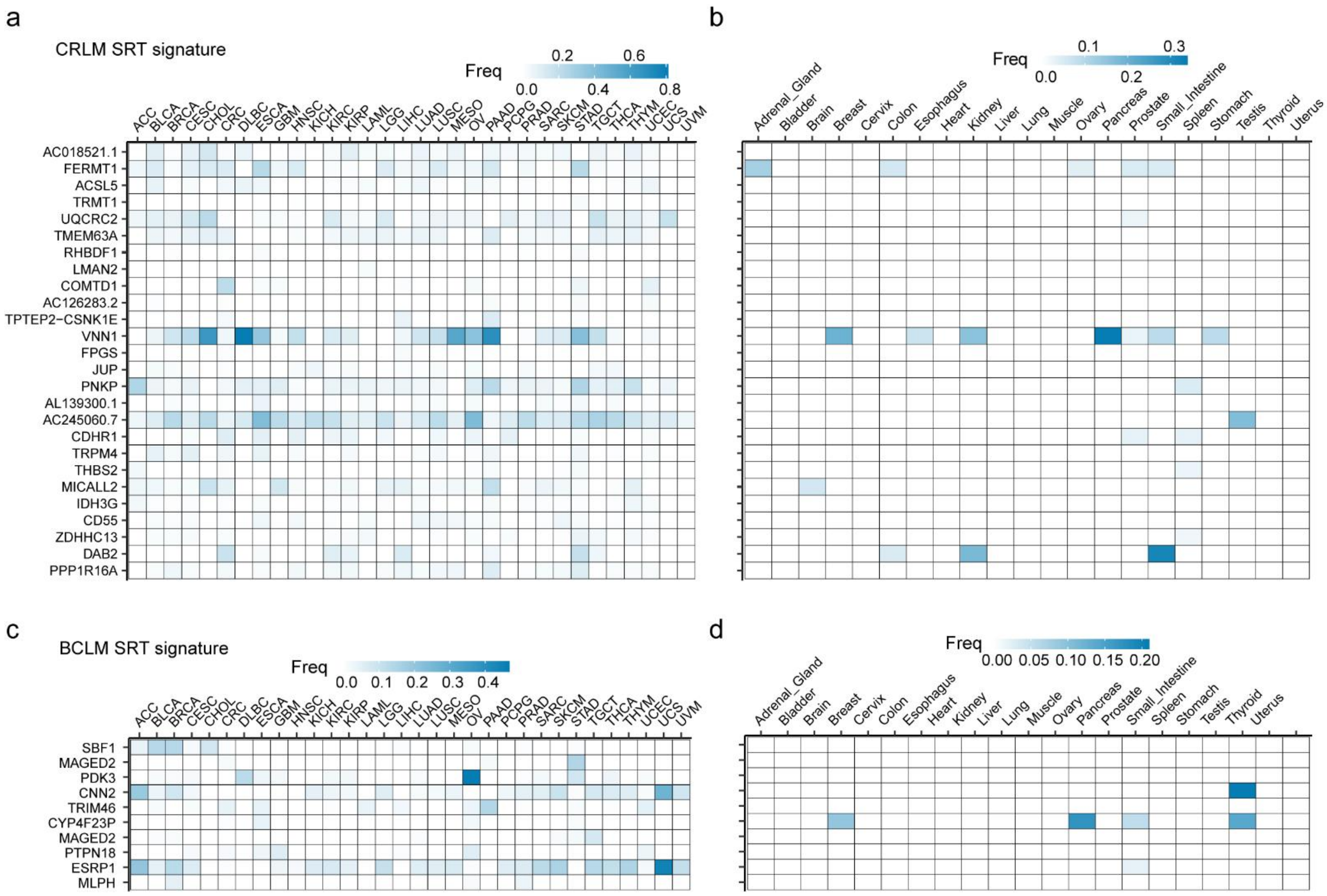


**Fig. S7 Metastasis-specific transcripts predict the metastasis and tissue origin of liver metastases**

a-b. Heatmap shows the expression of indicated CRLM-SRTs in tissues from the individual TCGA patients and normal tissues. c-d. Heatmap shows the expression of indicated BCLM-SRTs in tissues from the individual TCGA patients and normal tissues.

**Fig. S8**

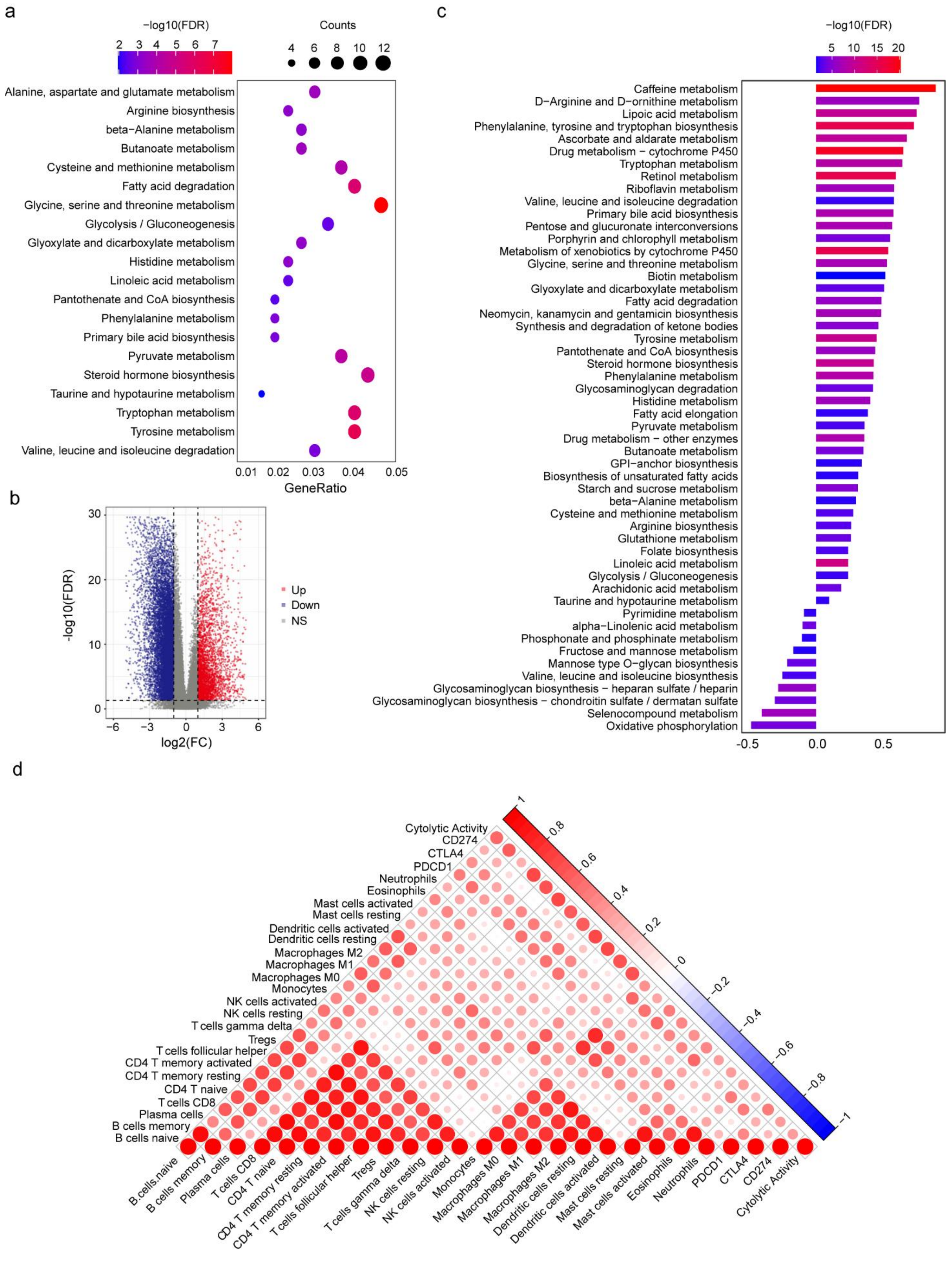
a
-log10(FDR)
2 3 4 5 6 7
Counts
4 6 8 10 12
Alanine, aspartate and glutamate metabolism
Arginine biosynthesis
beta−Alanine metabolism
Butanoate metabolism
Cysteine and methionine metabolism
Fatty acid degradation
Glycine, serine and threonine metabolism
Glycolysis / Gluconeogenesis
Glyoxylate and dicarboxylate metabolism
Histidine metabolism
Linoleic acid metabolism
Pantothenate and CoA biosynthesis
Phenylalanine metabolism
Primary bile acid biosynthesis
Pyruvate metabolism
Steroid hormone biosynthesis
Taurine and hypotaurine metabolism
Tryptophan metabolism
Tyrosine metabolism
Valine, leucine and isoleucine degradation
0.01 0.02 0.03 0.04 0.05
GeneRatio
b
-log10(FDR)
30 20 10 0
−6 −3 0 3 6
log2(FC)
Up
Down
NS
c
-log10(FDR)
5 10 15 20
Caffeine metabolism
D−Arginine and D−ornithine metabolism
Lipoic acid metabolism
Phenylalanine, tyrosine and tryptophan biosynthesis
Ascorbate and aldarate metabolism
Drug metabolism − cytochrome P450
Tryptophan metabolism
Retinol metabolism
Riboflavin metabolism
Valine, leucine and isoleucine degradation
Primary bile acid biosynthesis
Pentose and glucuronate interconversions
Porphyrin and chlorophyll metabolism
Metabolism of xenobiotics by cytochrome P450
Glycine, serine and threonine metabolism
Biotin metabolism
Glyoxylate and dicarboxylate metabolism
Fatty acid degradation
Neomycin, kanamycin and gentamicin biosynthesis
Synthesis and degradation of ketone bodies
Tyrosine metabolism
Pantothenate and CoA biosynthesis
Steroid hormone biosynthesis
Phenylalanine metabolism
Glycosaminoglycan degradation
Histidine metabolism
Fatty acid elongation
Pyruvate metabolism
Drug metabolism − other enzymes
Butanoate metabolism
GPI−anchor biosynthesis
Biosynthesis of unsaturated fatty acids
Starch and sucrose metabolism
beta−Alanine metabolism
Cysteine and methionine metabolism
Arginine biosynthesis
Glutathione metabolism
Folate biosynthesis
Linoleic acid metabolism
Glycolysis / Gluconeogenesis
Arachidonic acid metabolism
Taurine and hypotaurine metabolism
Pyrimidine metabolism
alpha−Linolenic acid metabolism
Phosphonate and phosphinate metabolism
Fructose and mannose metabolism
Mannose type O−glycan biosynthesis
Valine, leucine and isoleucine biosynthesis
Glycosaminoglycan biosynthesis − heparan sulfate / heparin
Glycosaminoglycan biosynthesis − chondroitin sulfate / dermatan sulfate
Selenocompound metabolism
Oxidative phosphorylation
-0.5 0.0 0.5
d
Cytolytic Activity
CD274
CTLA4
PDCD1
Neutrophils
Eosinophils
Mast cells activated
Mast cells resting
Dendritic cells activated
Dendritic cells resting
Macrophages M2
Macrophages M1
Macrophages M0
Monocytes
NK cells activated
NK cells resting
T cells gamma delta
Tregs
T cells follicular helper
CD4 T memory activated
CD4 T memory resting
CD4 T naive
T cells CD8
Plasma cells
B cells memory
B cells naive
B.cells.naive
1 0.8 0.6 0.4 0.2 0 −0.2 −0.4 −0.6 −0.8 −1

**Fig. S8. Altered transcriptome profiles and characteristics of the liver in metastatic liver cancer**

a. Pathways significantly enriched for genes with novel isoforms detected by Iso-Seq in non-tumor liver tissues from LM patients. b. Volcano plot of DETs in LM-NT compared to GTEx normal liver samples. Red and blue dots represent transcripts that were significantly upregulated and downregulated (FDR < 0.01), respectively. c. The difference in the relative activity of metabolism between LM-NT compared and GTEx normal liver samples. d. Correlations between tumor immunogenicity indicators, immune infiltration, and expression of immune checkpoint molecules.
